# Sequence programmable nucleic acid coacervates

**DOI:** 10.1101/2024.07.22.604687

**Authors:** Sumit Majumder, Sebastian Coupe, Nikta Fakhri, Ankur Jain

## Abstract

Nature uses bottom-up self-assembly to build structures with remarkable complexity and functionality. Understanding how molecular-scale interactions translate to macroscopic properties remains a major challenge and requires systems that effectively bridge these two scales. Here, we generate DNA and RNA liquids with exquisite programmability in their material properties. Nucleic acids are negatively charged, and in the presence of polycations, they may condense to a liquid-like state. Within these liquids, DNA and RNA retain sequence-specific hybridization abilities. We show that intermolecular hybridization in the condensed phase cross-links molecules and slows down chain dynamics. This reduced chain mobility is mirrored in the macroscopic properties of the condensates. Molecular diffusivity and material viscosity scale with the intermolecular hybridization energy, enabling precise sequence-based modulation of condensate properties over orders of magnitude. Our work offers a robust platform to create self-assembling programmable fluids and may help advance our understanding of liquid-like compartments in cells.

## Introduction

The sequence-specific binding properties of nucleic acids form the basis for life, providing the chemical foundation for the replication of genetic material, its decoding by the protein translation apparatus, and for molecular targeting such as by guiding RNA processing^1^. Nucleic acid hybridization is well-characterized both theoretically as well as experimentally^2–4^. This sequence-specificity of DNA bonding has been harnessed to generate a variety of self-assembling programmable materials^5^. These include nanometer-scale structures with precise control of DNA topology such as in DNA origami^6,7^ to mesoscale soft materials such as hydrogels^8,9^. In most of these applications, the negative charge on the phosphodiester backbone is neutralized by low valence counterions, and the desired molecular arrangement is achieved by the sequence-programmablity of DNA bonding.

Due to their charge, nucleic acids can also form electrostatic interaction-driven assemblies^10,11^. When solutions of oppositely charged polymers are mixed, they can undergo associative phase separation, called complex coacervation, resulting in a polymer-rich phase and a dilute supernatant phase^12,13^. DNA and RNA readily undergo complex coacervation in the presence of cationic polymers^11,14^. Complex coacervates concentrate nucleic acids without a membrane and this process may have parallels in the origin of life scenarios^15,16^. Eukaryotic cells also contain numerous membraneless organelles such as nucleoli and stress granules, and complex coacervation of RNA and proteins is implicated in the formation of these bodies^17–19^. Electrostatic interaction-mediated complexation is also the dominant approach used to assemble nucleic acids for delivering foreign genetic material to cells^20,21^.

Our previous work indicated that single-stranded DNA and RNA in complex coacervates retain their sequence-specific hybridization properties^22^. We hypothesized that we could harness this sequence specificity of nucleic acid hybridization in complex coacervates to generate new programmable materials. We were inspired by self-associative polymers that are designed to exhibit reversible associations between polymer chains^23,24^. These reversible inter-chain cross-links allow reconfiguration of the polymer network and manifest properties that are not accessible to conventional polymer materials with static cross-links, such as self-healing and stimuli-responsive material transformation^24^. These macroscopic properties depend on the lifetime, densities, and positions of the cross-links between polymer chains^25^. Precise engineering of these features in synthetic polymers is difficult, and the exquisite programmability of nucleic acid bonding could help overcome these challenges.

Here we generate DNA and RNA liquids with precise control over their viscoelastic properties. We engineered DNA oligonucleotides with short hybridization patches interspersed by single stranded regions (Fig. 1). Polyvalent cations induce complex coacervation resulting in DNA-rich liquids. Within the condensed phase, the DNA concentration is above the overlap concentration facilitating inter-chain hybridization. Such inter-chain hybridizations transiently cross link DNA molecules and slow down their diffusion. DNA mobility and the material properties of the condensed phase scale linearly with the life-time of DNA-DNA cross-links. By engineering the sequence, we can precisely program the hybridization energy and thus tune the properties of these liquids. Similar rules apply to condensates of RNA with cationic peptides. This system may pave the way for a new class of programmable nucleic acid-based materials. It may provide insights on the physics of polyelectrolyte complexation and may help uncover the rules underlying the formation of liquid-like RNA granules in the cell.

**Figure 1:**
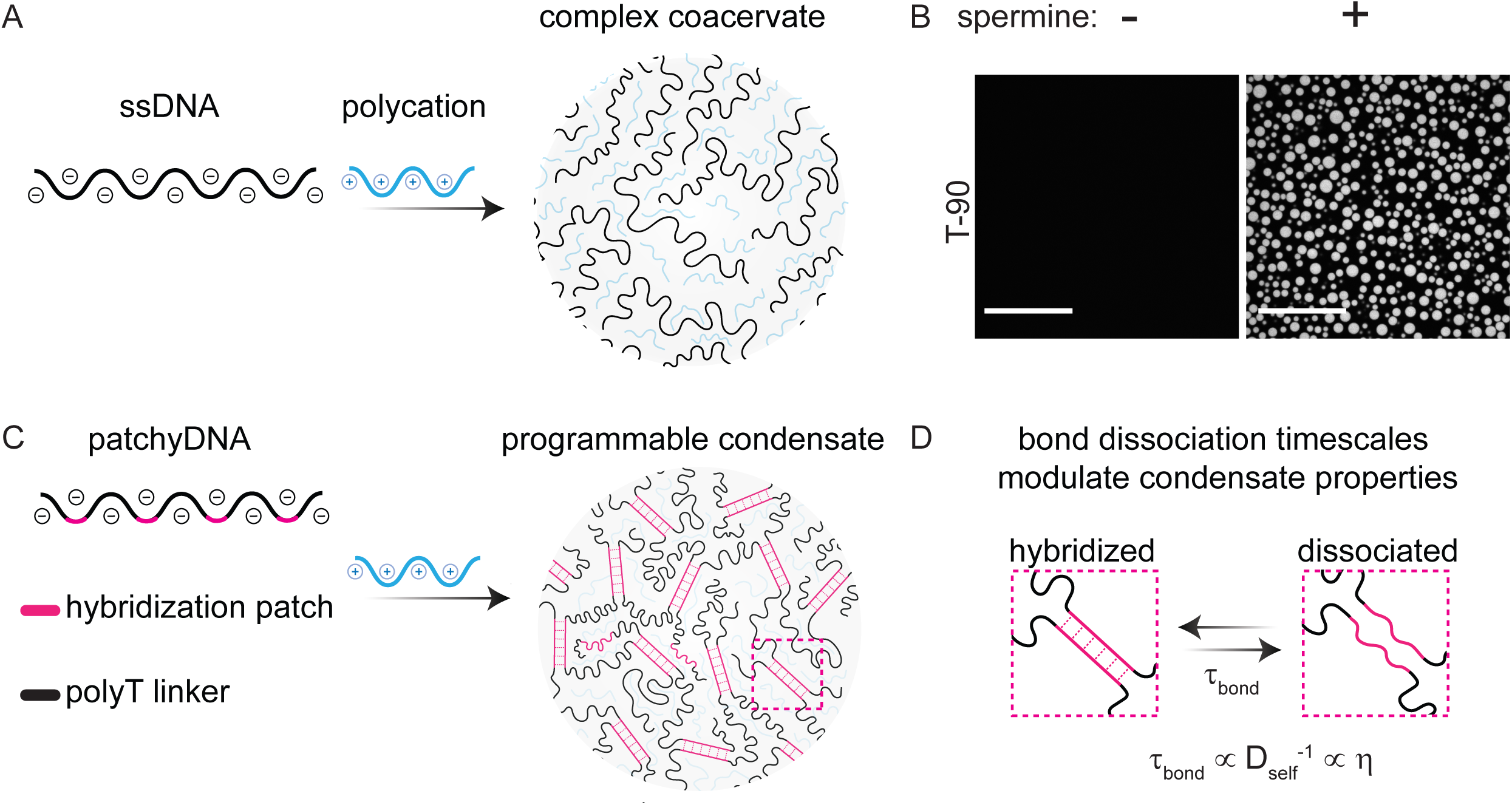
Scheme for generating condensates with sequence programmable properties. (A) Single stranded nucleic acids undergo associative phase separation in the presence of polyvalent cations and produce a polymer rich coacervate phase and a dilute supernatant phase. (B) Representative fluorescent micrographs showing complex coacervates of non-base pairing T-90 DNA in the presence of spermine. Scale bar, 30 µm. (C) We incorporated multiple weak hybridization patches in the DNA/RNA sequence. Inter-molecular hybridization at these sites transiently cross-links two or more strands and impedes their molecular mobility within coacervates. (D) Molecular diffusivity (D_self_) and viscosity (η) of these coacervates depend on the lifetime of inter-molecular cross-links (τ_bond_), and can be programmed by modulating the DNA sequence.

## Results

### Design of nucleic acids with self-associative patches

We used a 90-nucleotide long deoxythymidine (T-90) oligonucleotide as a chassis to build self-associative DNA (Fig. 1A-B, Fig. S1A). T-90 does not form appreciable secondary structure at room temperature and provides a benign scaffold to engineer base-pairing sites. To incorporate associative sites, we replaced a subset of T-bases in T-90 with short palindromic interaction patches (Fig. 1C-D, Table S1). These patches provide sites for intra- and inter-strand DNA hybridization while the total charge along the phosphodiester backbone is unchanged. The patch sequences were chosen such that the DNA transiently hybridize at room temperature (hybridization energy, ε = -ΔG/(k_b_TN_A_), between 0 and 20, where ΔG is the theoretical predicted free energy of hybridization from nearest neighbor calculations^26^, k_b_ is the Boltzmann constant, T is the temperature and N_A_ is the Avogadro constant, see Supplementary text S1 for additional notes on patch design). Successive patches were separated by ≥15 non-hybridizing T bases in order to minimize the cooperativity in hybridization between adjacent sites^27,28^. By changing the patch sequence or temperature, we can program the hybridization energy of the associative patch (Table S1 and S2) and varying the number of patches per DNA allows us to tune the hybridization valency. We refer to these self-associating single-stranded DNA oligonucleotides as patchyDNA drawing parallels to colloids with multiple interaction sites^29^.

As a polycation, we chose spermine, a cellular metabolite and a tetravalent cation at pH 7.0 (pKa for the four amine groups ≥ 7.9)^30^. Addition of spermine induced complex coacervation and produced two coexisting aqueous phases: a DNA-poor supernatant phase and a DNA-rich, condensed phase (Fig. 1B, Fig. S1). A small amount (< 1%) of fluorescently labeled DNA was doped to facilitate condensate visualization. The patchyDNA condensates exhibited liquid-like properties (Fig. S1B-E): they were spherical, and upon contact, two or more droplets coalesced and relaxed to a spherical geometry (Fig. S1E). Nearly 90% of DNA partitioned in the condensed phase (Fig. S2), and DNA partitioning and coacervate yield was comparable for the various sequences examined (Fig. S2).

Previous work has shown that the complex coacervate phase can be envisioned as a semi-dilute solution, i.e., within the condensed phase, the polymer concentration is comparable to the overlap concentration, making inter-chain interactions as likely as intra-chain interactions^31,32^. Consistent with this notion, the DNA concentration in the patchyDNA coacervates (C_dense_ ∼ 1-10 mM, Fig. S2) was comparable to the estimated overlap concentration (C_overlap_, for a 90-mer ssDNA ∼1 mM) (see Supporting text S2). We also used fluorescence resonance energy transfer (FRET) to directly estimate the mean inter-molecular distance between DNA in the coacervate and dilute phases. We labeled patchyDNA with either a fluorescence donor (Cy3) or an acceptor (Cy5) dye (Table S3). In the condensate, we observed substantial FRET between the two dyes (Fig. S3), indicating that the distance between DNA molecules is comparable to the molecular length scales (Forster radius for Cy3 and Cy5 dye pair ≈ 5 nm^33^; radius of gyration for 90-mer ssDNA, ≈ 5 nm^34^). In contrast, no measurable FRET signal was observed in the solution phase indicating that the hybridization patches do not induce stable DNA dimerization (Fig. S3).

As additional evidence for inter-molecular hybridization in the condensed phase, we examined the effect of base-pairing interactions on the phase stability. Complex coacervation is driven by a combination of attractive electrostatic interactions and entropically favorable molecular rearrangements. The loss in configurational entropy upon polyelectrolyte complexation is counterbalanced by the entropy gain upon the release of low-valence counter-ions that were bound to the polyelectrolytes^12,13,35^. High concentrations of low valence counter ions diminish the relative entropic gain upon polyelectrolyte complexation and inhibit complex coacervation^35^. Short-range intermolecular interactions such as hydrogen bonding and base stacking may stabilize complex coacervates against salt-mediated dissolution^36^. Consistent with this expectation, DNA-spermine coacervates were susceptible to NaCl mediated dissolution and the critical salt concentration for coacervate dissolution (C*) increased progressively with the patch hybridization energy (C* = 30 mM for ε = 0, 130 mM for ε = 20.2, Fig. S4). In summation, these results demonstrate that patchyDNA molecules are densely packed in complex coacervates and may form intermolecular base-pairs.

### Base-pairing modulates chain dynamics in patchyDNA condensates

Inter-molecular base-pairing would cross-link DNA strands and impede their mobility. To examine the effect of inter-strand cross-linking on chain dynamics, we used fluorescence recovery after photobleaching (FRAP). In these experiments, a small region of a DNA droplet is photobleached and the recovery is tracked over time. The fraction of fluorescence recovery reports on the proportion of DNA that is mobile within the droplet, while the recovery timescales reported on the molecular diffusivity. For the sequences examined (ε ≤ 20.2) (Fig. 2A, B), the patchyDNA coacervates exhibited near-complete (≥ 80%) fluorescence recovery upon partial photobleaching (Fig. 2C-D, Fig. S5A), indicating that the DNA in the condensed phase is mobile. However, the characteristic fluorescence recovery time, τ_FRAP_, progressively increased with ε, indicating that DNA mobility is reduced as the hybridization energy increases (Fig. 2C-D, Fig. S5A-B). τ_FRAP_ can be used to estimate the apparent self-diffusion coefficient, D_self_ (see Methods)^37,38^. D_self_ for the non-base-pairing T-90 was 0.32 ± 0.16 µm^2^/s (mean ± standard deviation, n = 16). As the hybridization energy of the patch was increased, D_self_ progressively decreased by nearly three orders of magnitude (D_self_ = (3.6 ± 1.3) × 10^−4^ µm^2^/s at ε = 20.2, mean ± standard deviation, n = 18, Fig. S5C). Base-pairing is temperature sensitive, and besides changing the DNA sequence, ε can also be modulated by changing the temperature (Table S2). For a given interaction patch, D_self_ could be modulated by nearly 5-fold as the temperature was increased from 20°C to 45°C (for patch sequence GGATCC, D_self_ = (2.6 ± 0. 3) × 10^−3^ µm^2^/s at 20°C, ε = 16.1, versus (17 ± 2) × 10^−3^ µm^2^/s at 45°C, ε = 10.8) (Fig. 2E, Fig. S6D). We did not observe any measurable change in D_self_ for T-90 with temperature, indicating that the DNA-spermine electrostatic interaction is not substantially affected in this temperature range (Fig. S6A and S7).

**Figure 2:**
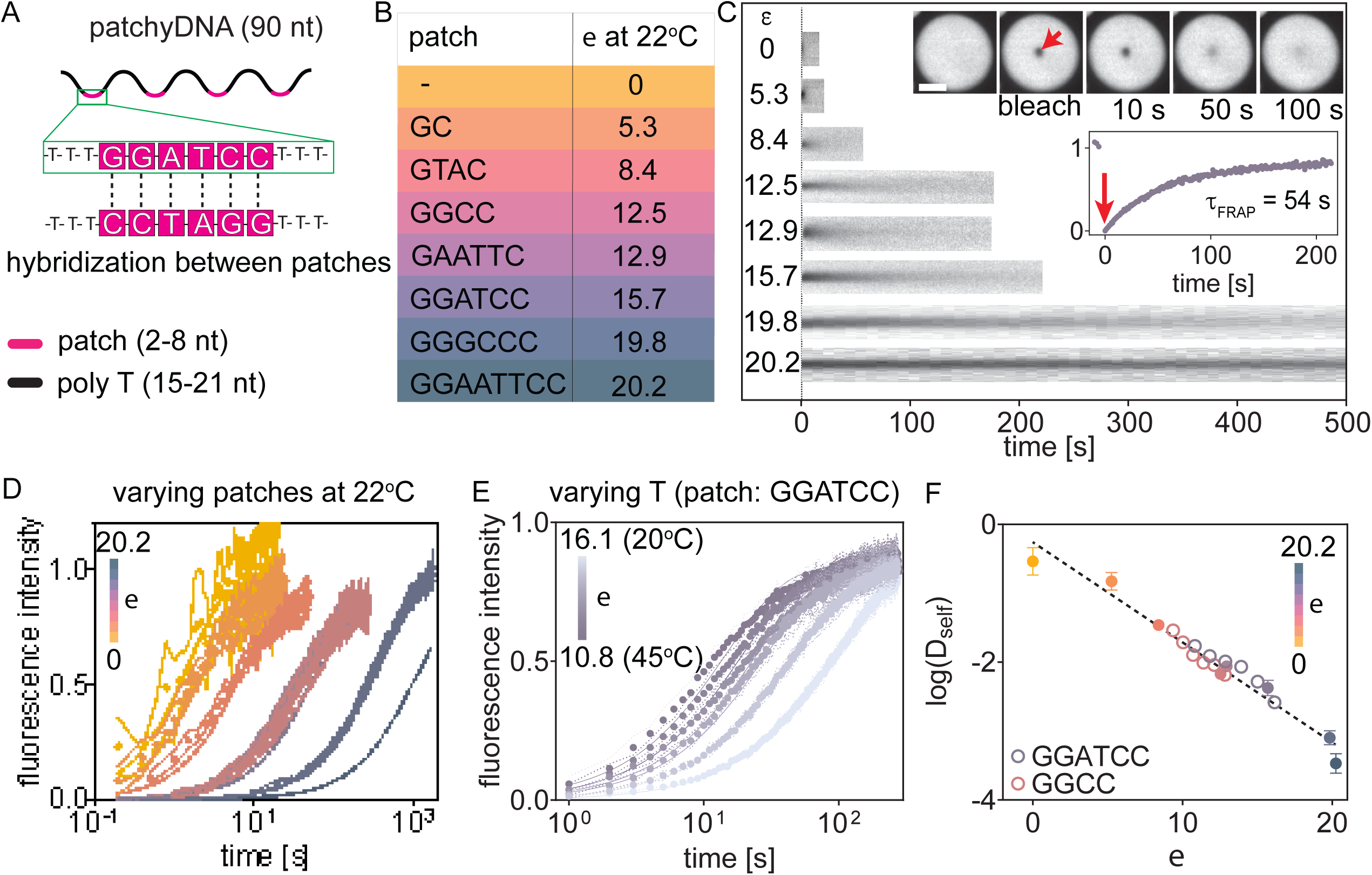
Base-pairing interactions modulate chain dynamics in patchyDNA condensates. (A) Schematic showing the design of self-associative patchyDNA. The hybridization patches are 2-8 nucleotide long palindromic sequences, and successive patches are separated by ≥15 T bases. (B) Representative patch sequences and the corresponding hybridization energies, ε, at 22^°^C. (C) Kymographs depicting fluorescence recovery upon partial photobleaching of patchyDNA condensates with the indicated ε. Inset: representative fluorescence micrographs and corresponding fluorescence time trace for DNA with the patch sequence GGATCC. The arrow indicates the point of photobleaching. (D, E) Fluorescence recovery time traces for patchyDNA condensates, where ε was changed either by varying the patch sequence (measurements conducted at 22°C) (D) or by varying the temperature for a given sequence (E). Each data point denotes mean ± SD, n ≥ 5 droplets. (F) Plot showing that logarithm of DNA self-diffusion coefficient (D_self_) scales linearly with ε. The dotted line is the least square fit between log(D_self_) and ε (slope = −0.14 ± 0.01, R^2^ = 0.90). The solid and open symbols in (F) denote that ε was varied by changing the patch sequence or by changing the temperature respectively. The color bars in D, E and F denote the range of ε. Scale bar in C is 5 µm.

Interestingly, all of our D_self_ measurements across various patch sequences and temperatures, when plotted against ε, converged on a single line on the semi-log plot, indicating that D_self_ scales exponentially with ε (Fig. 2F). This exponential scaling between D_self_ and ε is reminiscent of the dynamics of unentangled self-associative neutral polymers in the semi-dilute regime^25,39^. According to this model (referred to as the sticky Rouse model), associative polymers in a network diffuse by breaking inter-molecular cross-links on one site and re-forming at another^25,40^. Thus, polymer diffusion is inversely related to the lifetime of intermolecular cross-links, τ_bond_^39,40^. The bond lifetime is determined by the interaction energy and is given by τ_bond_ ∼ e^−Ea/kbT^, where E_a_ is the association energy, and k_b_T is the normalization for thermal energy. While we do not know the exact solvent environment in the condensed phase, these results indicate that the theoretically estimated hybridization energy provides a reasonable proxy for the association energy and demonstrate that DNA diffusion in the coacervate phase can be precisely programmed by engineering the hybridization patch sequence.

### Inter-chain hybridization modulates material properties of DNA coacervates

We next examined the viscoelastic properties of coacervates using particle tracking microrheology^41^. In these experiments, fluorescent microspheres are embedded in the material and their movement over time reports on the material’s viscoelastic behavior^41,42^. Beads trapped in the DNA coacervates exhibited random fluctuations that are effectively described by a Gaussian probability distribution (Fig. 3A-B, Fig. S8). The mean squared displacement (MSD) of beads followed a power law relation with time (t), MSD ∼ t^α^, with diffusion exponent α ≈ 1, across the various patch sequences examined (Fig. 3C-D). This linear relationship between bead MSD and time indicates that the patchyDNA coacervates behave like viscous fluids with minimal elastic behavior and aligns with our observations from fluorescence photobleaching experiments described above.

**Figure 3:**
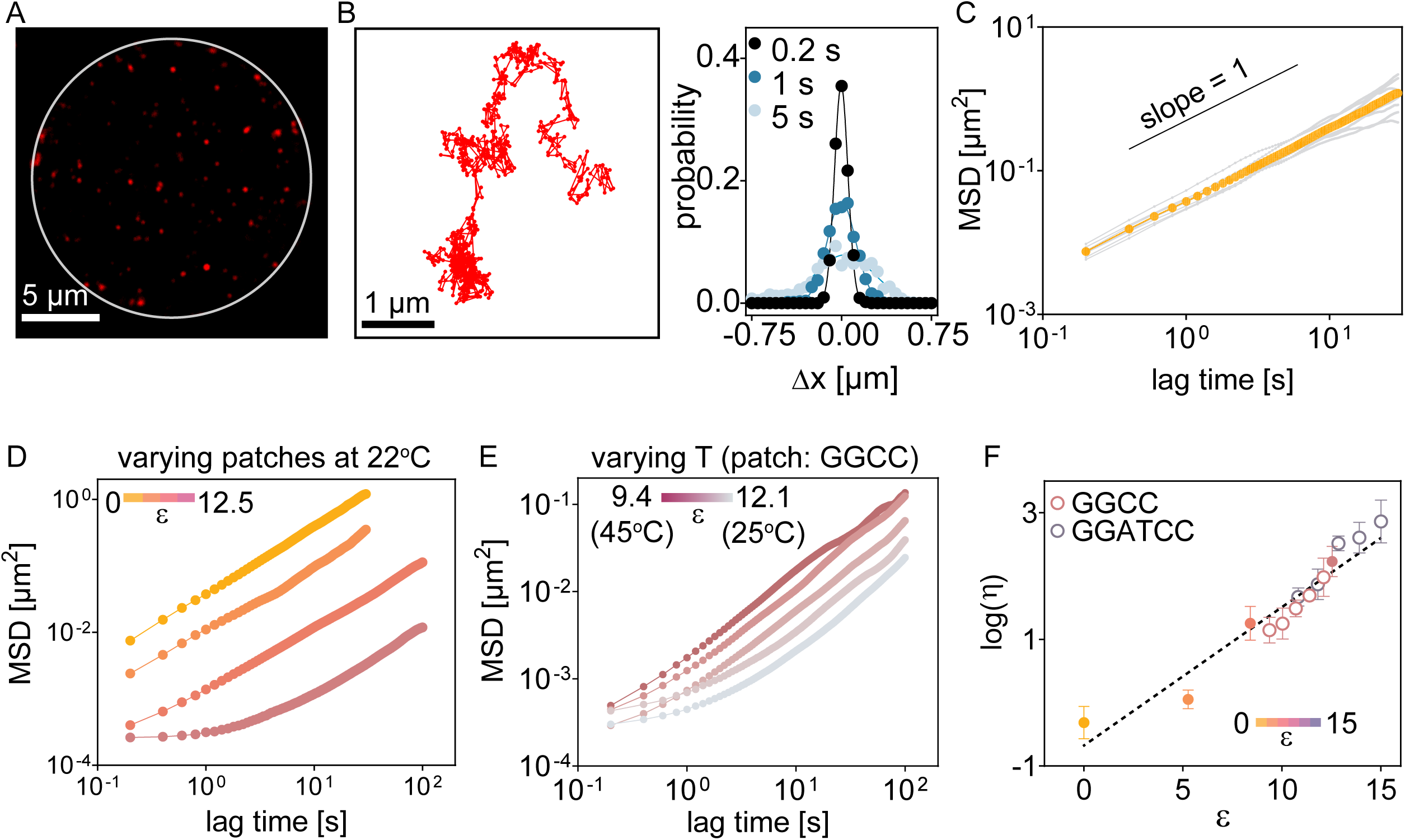
Base-pairing interactions modulate viscosity of patchyDNA condensates. (A) Representative fluorescence micrograph showing passivated beads (radius = 50 nm) embedded in a patchyDNA coacervate (white circle). (B) Representative time trajectory of a single bead (left) and probability distribution of bead displacement at three different lag times (right) in T-90 coacervates at 22^°^C. Distributions are well-described by a Gaussian function (solid lines). (C) Mean squared displacement (MSD) with lag time for beads embedded in T-90 condensates at 22^°^C. Grey lines show the data from individual beads. Mean values are in yellow. Black solid line is provided as a guide for slope = 1. (D) MSD tracks of beads (n > 5, mean) in various patchyDNA condensates at 22^°^C. (E) Similar to (D) but for beads in GGCC coacervates at different temperatures. (F) Plot showing the logarithm of viscosity (*η*) versus ε. The solid and open symbols denote ε was varied by changing the patch sequence or by changing the temperature respectively. The dotted line is a linear fit between log(*η*) and ε (slope = 0.22 ± 0.01, R^2^ = 0.88). The color bars in D, E and F denote the range of ε.

The bead fluctuations can be used to calculate its diffusion coefficient, D_probe_, using the relationship, MSD ≈ 4·D_probe_ t + NF, where NF is the noise floor (see Methods, Fig. S9). Since these coacervates behave like viscous liquids and knowing the size of the bead (see Methods), we can use Stokes-Einstein relation to estimate the material’s dynamic viscosity (*η*). *η* increased with ε, increasing from 0.56 ± 0.32 Pa.s (for T-90, ε = 0; mean ± standard deviation, n = 7) to 191 ± 97 Pa.s (for patch sequence GGCC, ε = 12.53; mean ± standard deviation, n = 6). Likewise, for a given DNA sequence, *η* increased when ε was increased by varying the temperature (Fig. 3E, Fig. S10). Similar to our observations with D_self_, *η* scaled exponentially with ε (Fig. 3F), indicating that the macroscopic properties of the condensates can be programmed by the DNA sequence. We limited our microrheology experiments to DNA with ε ≈ 15 as beyond this regime, the bead diffusion approached the measurement noise floor (Fig. S9).

When two liquid droplets fuse, interfacial tension drives them to minimize the surface area by relaxing to a spherical shape, while viscosity opposes this relaxation. For droplets of a viscous fluid suspended in a lower viscosity medium, the timescales of fusion (τ_fusion_) are proportional to the droplet viscosity (*η*) and its size (*l*), and inversely proportional to the interfacial tension (*γ*)^43,44^. Consistent with their liquid-like behavior, patchyDNA droplets fused with one another (Fig. 4A). For a given patchyDNA sequence, τ_fusion_ increased linearly with the droplet size, *l*, and the slope of this line provided an estimate of the ratio, *η*/*γ*, also knowns as the inverse capillary velocity (Fig. 4B-D and Fig. S11A). We estimated *η*/*γ* for the various patchyDNA sequences by examining multiple coalescence events across a range of droplet sizes and found that *η*/*γ* increased by 6 orders of magnitude as ε was varied from 0 to 20.2 (Fig. 4E). The droplet coalescence rates, in conjunction with direct viscosity measurements, allowed us to estimate the droplet interfacial tension. Interfacial tension for patchyDNA coacervates was between 400 −1200 µN/m, about two orders of magnitude smaller than that for air-water interface at 22^°^C (≈ 70 mN/m)^45^ (Fig. S11B). This ultra-low interfacial tension is comparable to that for condensates of other polymers of similar molecular dimensions^46,47^. Within the constraints of our experiments, *γ* did not appreciably change with ε (Fig. S11B). The fusion experiments allowed us to extract *η*/*γ* for a larger spectrum of patchyDNA, and in conjunction with our observation that *γ* exhibits a weak dependence on ε, these results indicate that *η* scales exponentially with ε, and can be modulated over 6 orders of magnitude as ε varies from 0 to 20.2 (*η*/*γ* ≈ 5.2 × 10^−4^ sµm^−1^ for ε = 0, ≈ 2.9 × 10^2^ sµm^−1^ for ε = 20.2). Altogether, our data demonstrate that by programming the sequence of the DNA patch, we can tune the life-time of the DNA-DNA cross-link in complex coacervates and this interchain interaction is reflected in the molecular diffusion and macroscopic properties of the condensed phase.

**Figure 4:**
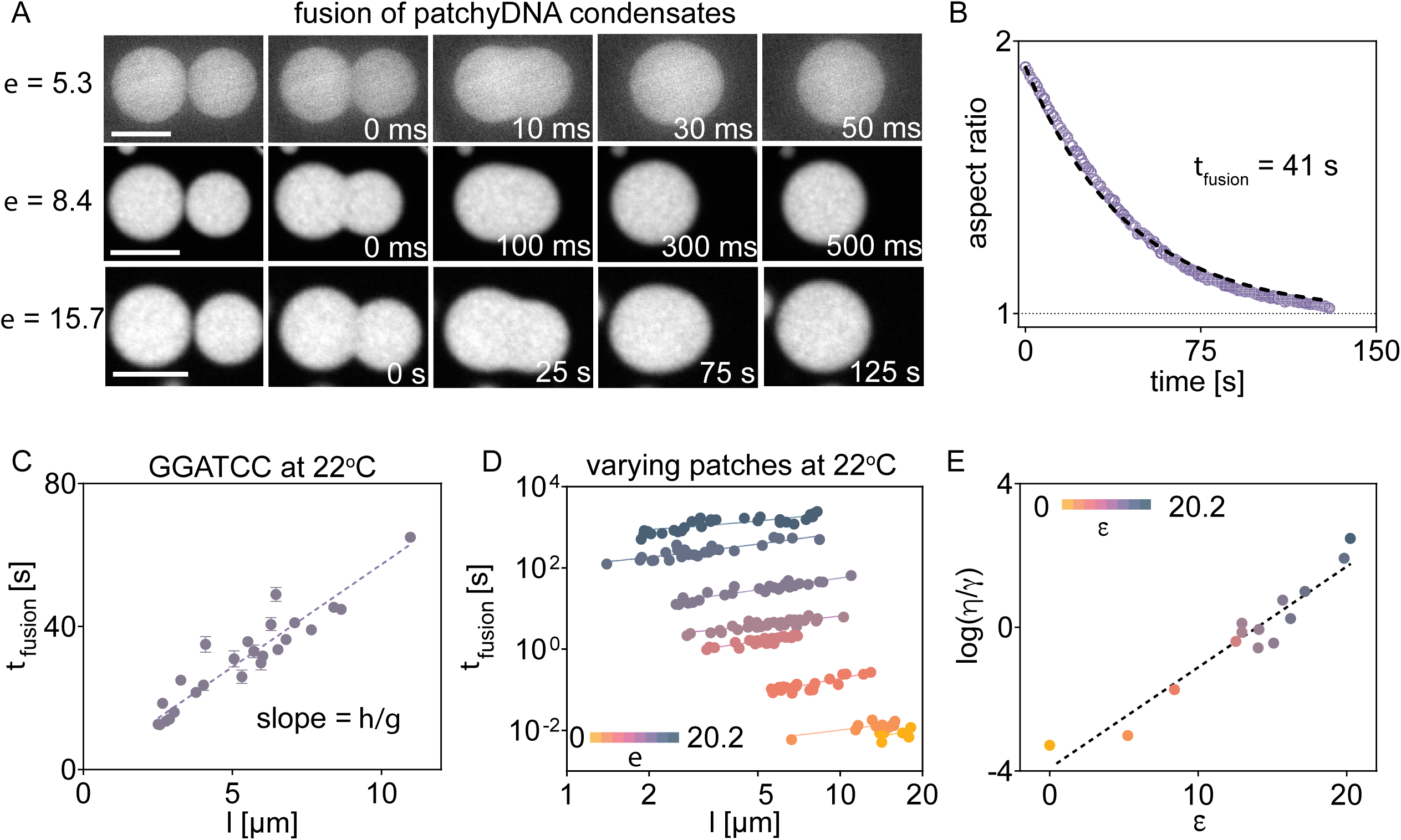
Base-pairing tunes the fusion dynamics of patchyDNA condensates. (A) Representative fluorescence micrographs showing fusion between complex coacervates of patchyDNAs with the indicated ε at 22^°^C. Fusion rates are slower for DNA with higher ε. Scale bar, 10 µm. (B) The characteristic fusion time (τ_fusion_) can be obtained from fitting single exponential (dotted line) to the aspect ratio of the fusing droplets of roughly similar size (*l*). The plot in (B) corresponds to ε = 15.7 in (A). (C) τ_fusion_ linearly increased with the droplet size (*l*). Representative data from patchyDNA coacervates with patch sequence GGATCC, ε = 15.7. (D) τ_fusion_ for various patchyDNA coacervates increases linearly (solid lines) with *l*. The color bar indicates ε at 22^°^C. (E) Plot showing that the logarithm of inverse capillary velocity (*η*/*γ*) scales linearly with ε (dotted line, slope = 0.28 ± 0.02, R^2^ = 0.92). The color bars in D and E show the corresponding range of ε.

### Heterotypic interactions enable design of stimuli-responsive condensates

The DNA in the coacervates can also be potentially cross-linked via heterotypic interactions. Such heterotypic associations could allow one to tune the coacervate properties in response to a trigger strand that potentiates inter-molecular cross-linking (Fig. 5A, S12A). As a proof of concept, we designed two sets of patchyDNAs: a primary strand and an actuator secondary strand (Fig. 5A). The patches on these DNAs individually do not self-associate, but they can hybridize to each other (Fig. 5A-B). The coacervates produced from the primary strand alone (patch sequence, CTCCTC) exhibited dynamics and material properties similar to those from T-90 DNA, as expected (Fig. S12B). Upon addition of the cross-linking secondary DNA (patch sequence, GAGGAG), the self-diffusivity of the primary strand diminished (Fig. S12C). The apparent self-diffusion coefficient, D_self_, was lowest when the two strands were mixed at equimolar ratio and increased when either strand was in excess (Fig. S12C).

**Figure 5:**
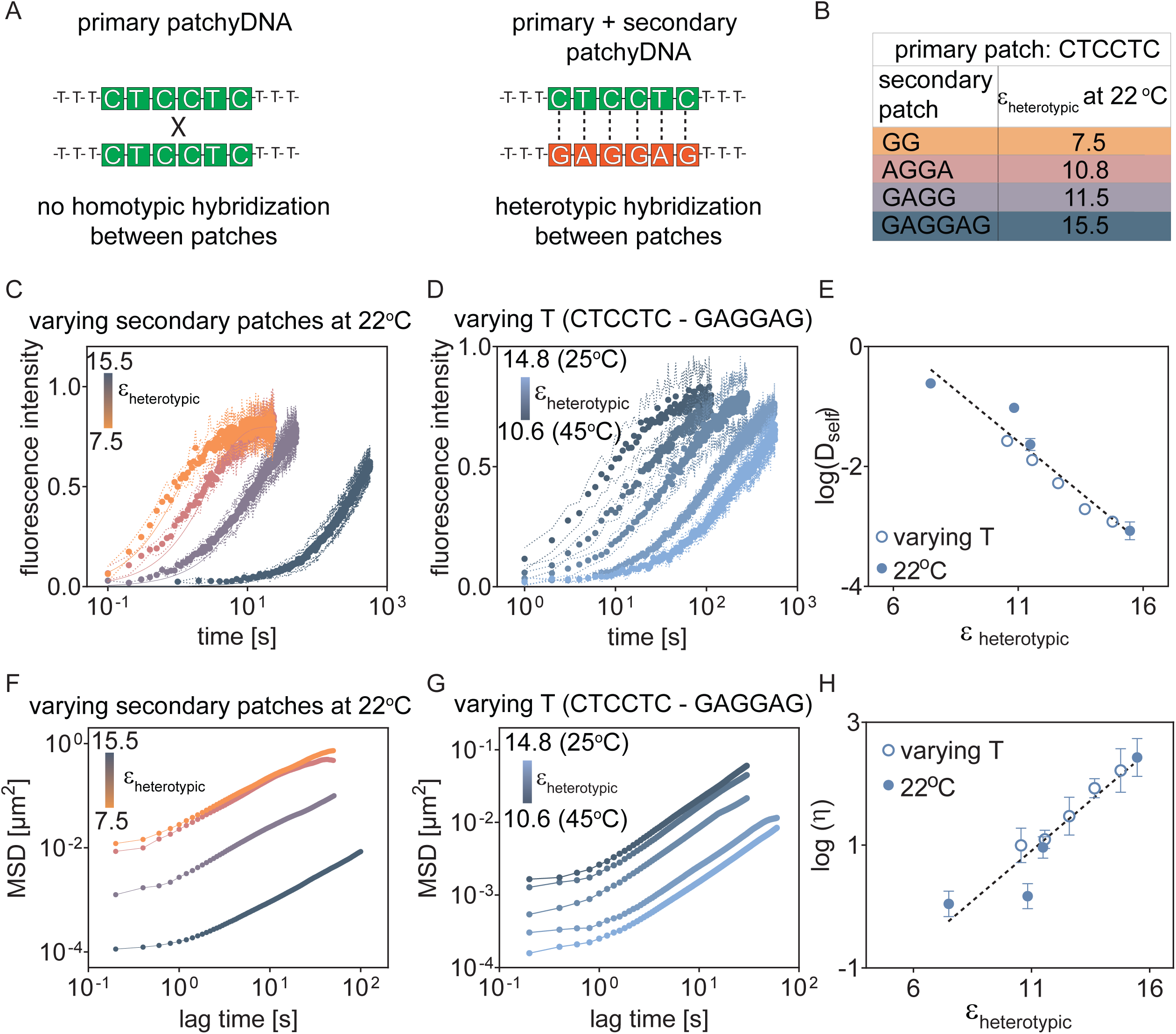
Heterotypic base-pairing modulates molecular dynamics and viscosity of patchyDNA condensates. (A) Schematic showing design of a primary patchyDNA with inducible inter-molecular cross-linking. The primary strand forms complex coacervates with spermine but this sequence does not contain self-associative hybridization sites (left). Addition of an appropriate secondary patchyDNA strand may cross-link the two DNAs (right). (B) Estimated hybridization energy for heterotypic cross-links (ε_heterotypic_) with the indicated secondary patch sequence. Patch sequence for primary strand is CTCCTC. (C, D) Fluorescence recovery after photobleaching for the primary patchyDNA and (F, G) MSD of embedded beads with lag time, where ε_heterotypic_ was modulated either by varying the secondary patchyDNA (C, F) or by varying the temperature for the indicated secondary patchyDNA (D, G). Data depict mean ± SD, n ≥ 5 and mean, n ≥ 5 for (C, D) and (F, G), respectively. (E, H) Plots showing that the logarithm of self-diffusivity (E) and viscosity (H) of the primary patchyDNA scales linearly with ε_heterotypic_. Dotted line denotes the least square fit (slope = −0.34 ± 0.01, R^2^ = 0.92 and 0.33 ± 0.02, R^2^ = 0.83, for E and H respectively). The solid and open symbols indicate that ε_heterotypic_ was varied by changing the secondary patchyDNA or by changing the temperature, respectively. The color bars in C, D, F, G indicate the range of ε_heterotypic_.

This ability to harness heterotypic interactions allowed us to tune the properties of the coacervates produced from the same primary patchyDNA but in response to distinct cross-linking secondary DNAs as triggers. We examined four distinct secondary sequences that could each hybridize with the same primary patchyDNA but with different hybridization energies (Fig. 5B). Similar to our observations with self-associative patches, both D_self_ and *η* scaled exponentially with the inter-molecular hybridization energy, ε_heterotypic_, when the two strands were mixed at stoichiometric ratios (Fig. 5C-H). These results, in conjunction with the findings from previous sections, reinforce that the material properties of DNA complex coacervates can be predictably tuned by modulating hybridization-based cross-links, including in response to an external input.

### Inter-molecular hybridization modulates the material properties of RNA-peptide condensates

Eukaryotic cells are compartmentalized by numerous RNA-containing condensates such as nucleoli, nuclear speckles, and stress granules, that exhibit liquid-like properties^17,48^. The protein constituents of these condensates often contain charged disordered domains^49^. In vitro, many of these proteins forms condensates in the presence of RNA^18,50^. The choice of RNA sequence affects condensate properties^37,51^ but the role of RNA-RNA interactions in these bodies remains poorly characterized^52^. Equipped with our biophysical framework, we examined how inter-molecular RNA-RNA hybridization affects the properties of RNA condensates produced with cationic peptides.

We used a 60-nucleotide long polyU (rU-60) RNA and incorporated 4 associative patches that are expected to reversibly hybridize at room temperature (Fig. 6A). rU-60 does not form appreciable secondary structure at or above room temperature (22°C)^11^. As a cationic peptide, we used poly-l-lysine (K10). The pKa of the lysine side chain is 10.5, and this peptide is expected to be fully protonated at the neutral pH^53^. The K10 peptide induced RNA complex coacervation (Fig. 6A). These coacervates exhibited liquid-like behavior and upon fusion of two or more droplets, they rapidly relaxed to a spherical geometry (Fig. S13A). The fusion timescales, τ_fusion_, increased linearly with the droplet size, indicating that these coacervates behave like simple viscous liquids (Fig. S13A). The measured viscosity, *η*, for rU-60/K10 complex coacervates was 4.4 ± 1.6 Pa.s (mean ± standard deviation, n = 5) and increased progressively by about three orders of magnitude to 476 ± 96 Pa.s (mean ± standard deviation, n = 5) as the hybridization energy, ε_RNA_, was increased to 9.5, by either modulating the patch sequence (Fig. 6B) or by changing temperature (Fig. 6C, Fig. S14). The interfacial tension for RNA-peptide coacervates was ∼1 mN/s and only modestly varied with ε_RNA_ (Fig. S14B). Like patchyDNA, all measured values of *η* and *η/γ* across various perturbations could be described by a single exponential relationship (Fig. 6D-E). This exponential scaling between *η* and ε_RNA_ demonstrates that the dynamics of these droplets scale with the lifetime of RNA-RNA bonds and reinforces that the macroscopic properties of these RNA-containing condensates are tuned by inter-strand RNA hybridization.

**Figure 6:**
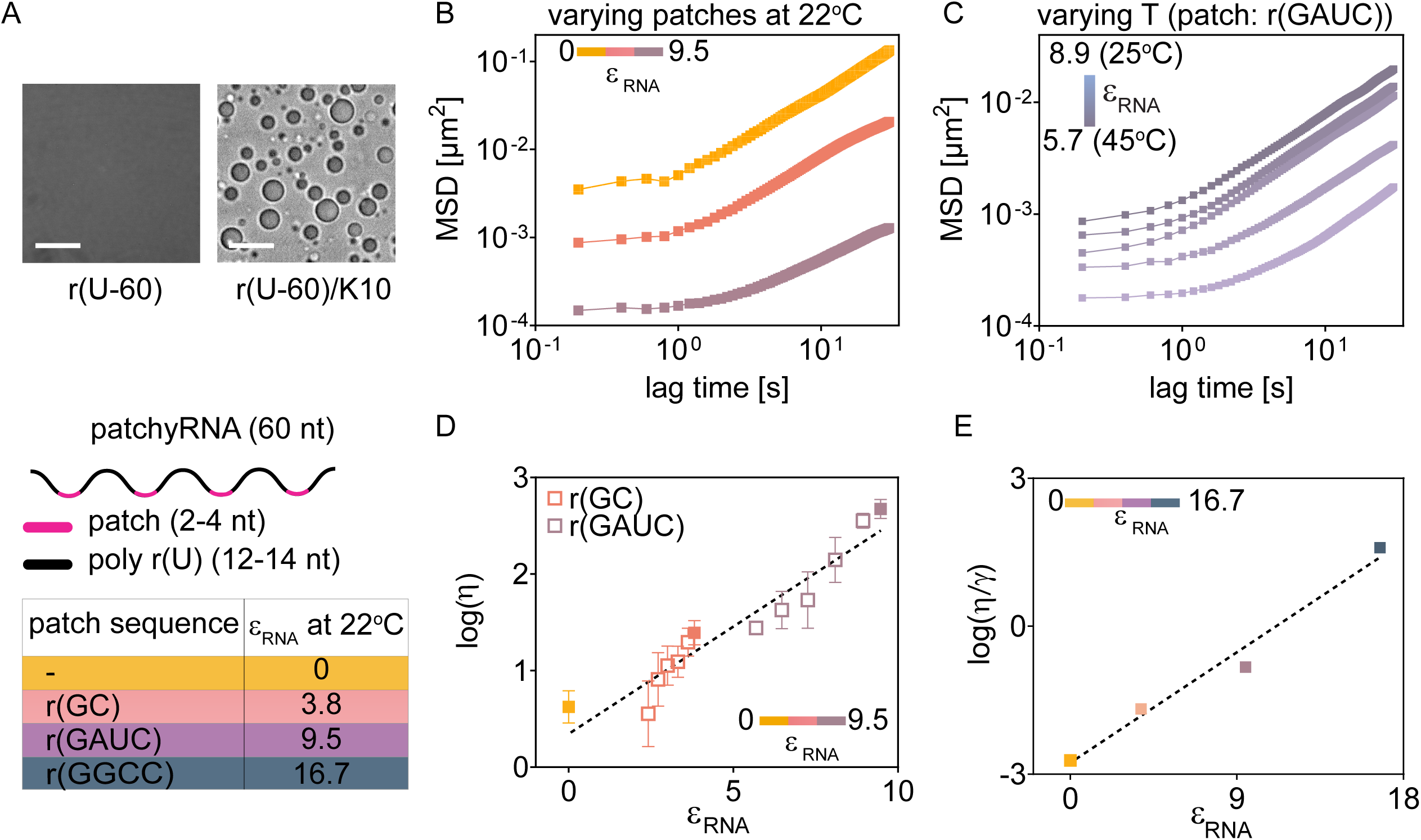
Inter-molecular base-pairing modulates the properties of RNA-peptide condensates. (A) Top: representative micrographs showing that non-base-pairing U-60 RNA forms complex coacervates with poly-l-lysine (K10). Scale bar, 10 µm. Bottom: schematic depicting the design of a 60-nucleotide long patchyRNA with four hybridization patches. Table shows hybridization energy of patches (ε_RNA_) at 22^°^C. (B, C) MSD of passivated beads in patchyRNA/K10 condensates, where ε_RNA_ was modulated either by varying the patch sequence (B) or by temperature (C). The color bars represent the range of ε_RNA_. Each datapoint is mean of n ≥5 beads. (D, E) The logarithm of viscosity (*η*) (D), and inverse-capillary velocity (*η*/*γ*) (E) scales linearly with ε_RNA_ (dotted line, slope = −0.22 ± 0.01, R^2^ = 0.85 and 0.25 ± 0.03, R^2^ = 0.98, respectively). The solid and open symbols in (D) denote ε_RNA_ was varied by changing the patch sequence or by temperature respectively.

## Discussion

We harnessed the two fundamental properties of nucleic acids, the negative charge and the sequence programmability of their bonding, to generate liquid-like materials with exquisite control over their physical properties. Nucleic acids in complex coacervates form electrostatic interaction mediated network, and super-imposing base-pairing interactions allowed us to modulate the chain dynamics. By engineering the sequence, we can program the lifetime of inter-strand cross-links, and these molecular scale interactions are reflected in the macroscopic properties of the condensates. We demonstrate programmability over orders of magnitude in chain mobility and condensate viscosity. The material properties scale exponentially with the predicted hybridization energy and allow us to exert striking control over the bulk properties by manipulating the sequence. This ability to tune material properties of DNA and RNA complex coacervates, in conjunction with their inherent stability and biocompatibility, may inspire a new class of nucleic acid-based self-assembling materials.

Polymer networks with dynamic and reversible associations exhibit properties that are not achieved in conventional polymers with static cross-links, and offer new functionalities such as ease of degradation, responsiveness to stimuli, and self-healing^23,24^. Given these desirable properties, there is considerable interest in generating new polymers with reversible associations, but it remains challenging to precisely engineer the strength and positioning of these interactions in synthetic polymers. Employing DNA hybridization for inter-chain interactions provides exceptional control over the engineerability of these association sites. Advances in DNA synthesis scales^54^ and development of chimeric DNA-conjugated polymers^55^ may provide a route to produce bulk-scale materials with precise control over material properties. There is also tremendous interest in understanding the physics of associative polymers, but limitations in the production of synthetic polymers with desired molecular patterns, interactions, and lengths, have hampered experimental examination. Our system may provide a facile platform to overcome these challenges and advance our fundamental understanding of associative polymer systems.

Our work also has implications towards understanding the assembly of biomolecular condensates in the cell. RNA provides scaffold for the formation of numerous biomolecular condensates but how the sequence-specific physicochemical properties of RNA affect its condensation is only beginning to be uncovered^52^. Previous works have shown that single-stranded and double-stranded nucleic acids, that differ in stiffness and charge density, yield coacervates with distinct properties^10,51^. We show that besides stable duplexes, transient sequence-specific inter-molecular base-pairing between RNA may profoundly alter condensate properties. These transient interactions impede molecular diffusion and may tune reaction rates. Analogous to the molecular grammar of condensate-associated proteins^56^, our work may lay the foundation for uncovering the RNA grammar in condensates. Mutations in condensate-associated proteins that alter the viscoelastic properties of condensates are observed in neurodegenerative diseases^57^. For example, ALS-causative mutations in FUS and TDP-43 reduce stress granules dynamics^58^. Our work shows that like proteins, RNA sequence and base-pairing interactions can have a profound effect on condensate properties and may potentially harbor disease causative mutations. In particular, GC-rich repeptitive sequences (such as tandem CAG and GGGGCC repeats) that are associated with neurodegenerative disorders produce elastic coacervates with minimal RNA mobility^22^. Our synthetic nucleic acid coacervate system may help uncover the principles that govern the formation and function of RNA granules in the cell and may be potentially used to create synthetic membraneless organelles in cells.

## Methods

### M1. Chemicals

Single-stranded DNA and RNA oligonucleotides were purchased from IDT, USA. Spermine tetrahydrochloride and poly-l-lysine hydrochloride (MW 1600 Da) were purchased from Millipore Sigma, USA (S2876), and Alamanda Polymers, USA (26124-78-7), respectively. Nuclease-free water (AM9939) and Tris buffers (pH 7.0, AM9850G and pH 8.0, AM9855G) were obtained from Thermo Fisher Scientific, USA. IDTE pH 7.5 was purchased from IDT, USA (11-05-01-15). DNA was resuspended at a concentration of 250 µM in TE buffer (10 mM Tris, pH 8.0, 0.1 mM EDTA). RNA was resuspended at a concentration of 200 µM in IDTE pH 7.5. Stock solutions of spermine hydrochloride (100 mM) and poly-l-lysine hydrochloride (25 mM) were prepared in nuclease-free water. Stock solutions were aliquoted and stored at −20°C and thawed immediately before use.

### M2. In vitro complex coacervation

DNA complex coacervates were prepared by sequentially adding the reagents to a PCR tube in the following order to the indicated concentrations: nuclease-free water, Tris buffer (pH 7.0, 10 mM), DNA solution (10 µM), labeled DNA (40 nM), spermine (4 mM), and NaCl (0–160 mM). The components were thoroughly mixed by pipetting. Immediately after mixing, the solution became turbid indicative of phase separation. The mixture was transferred onto a passivated glass surface for visualization under a microscope. For examining heterotypic interactions, the stock solutions of two DNA species were first mixed at the desired stoichiometric ratio, and complex coacervation was induced as described above. The resulting complex coacervates of DNA mixtures were heat denatured at 70°C for 3 minutes, and cooled down to room temperature before characterization. RNA coacervates were prepared by mixing RNA (16 µM) and poly-l-lysine (0.5 mM) in 10 mM Tris (pH 7.0).

### M3. Glass passivation

Glass-bottom 384-well plates (Brooks, MatriPlate MGB101-1-2-LG-L) and cover glass (VWR, USA, 48393-251) were passivated with PEG-silane (MW = 5 kD, Laysan bio, MPEG-SIL-5000-1g). Briefly, the glass surface was first activated with a 0.5 M NaOH (Millipore Sigma, USA, 1310-73-2) solution for 20 minutes at 40°C. This solution was removed, and the glass surface was rinsed three times using Milli-Q water and dried at 40°C for an additional 30 minutes. Subsequently, the glass surface was treated with 5% (w/v) PEG-silane in anhydrous ethanol and incubated at room temperature (22°C) for 20 minutes. After removing the PEG-silane solution, the glass surface was washed three times with Milli-Q water and dried using a gentle flow of dry air. The passivated glasses were used immediately or stored at 4°C.

### M4. Microsphere passivation

Red-fluorescent carboxy-coated polystyrene beads (diameter = 100 nm, Invitrogen^TM^ FluoSpheres^TM^ F8801) were passivated with amine-terminated methoxy-PEG (mPEG-NH2; MW 750 Da; Millipore Sigma 07964) following the protocol described by Valentine, M. T. et al. (2004)^59^. Briefly, beads were diluted to 4 × 10^12^ particles/ml and tip-sonicated (10% amplitude, 5 seconds on, 5 seconds off, 30 seconds total time) to minimize inter-bead adhesion. The diluted bead solution was dialyzed into 100 mM 2-(N-morpholino)ethanesulfonic acid (MES, 150x volume, Millipore Sigma M3671) at pH 6.0 for 2.5 hours. The dialysis bag was rinsed with deionized water and transferred to a solution mixture of 100 mM MES, 15 mM 1-[3-(dimethylamino)propyl]-3-ethylcarbodiimide (Millipore Sigma E6383), 5 mM N-hydroxysuccinimide (NHS) (Millipore Sigma 130672), and a 10x excess of mPEG-NH2. After 40 minutes of reaction, the dialysis bag was transferred to borate buffer (50 mM boric acid, Millipore Sigma B0394, 36 mM sodium tetraborate, Millipore Sigma 221732) at pH 8.5 containing 5 mM NHS and a 10x excess of mPEG-NH2 and dialyzed for at least 8 hours. This dialysis protocol was repeated two more times before transferring the dialysis cassette to pure borate buffer, where dialysis continued for another 4 hours. Upon completion of dialysis, the bead solution was removed from the dialysis cassette, diluted (50x volume) into 10 mM Tris (Millipore Sigma 93352) pH 7.0 buffer, and stored at 4 °C for future use.

### M5. Microscopy

Complex coacervates were visualized using an Andor Dragonfly 500 spinning disk confocal system (Oxford Instruments, USA) mounted on a Nikon TE2000E inverted microscope using a 100x Nikon oil immersion objective (NA 1.45). Images were acquired an Andor iXon EM-CCD or an Andor Zyla sCMOS (Oxford Instruments, USA). Images were captured using 561 nm (for Cy3 labeled DNA), and 637 nm (for Cy5 labeled DNA) excitation lasers. The imaging scan speed was varied between 200 and 0.1 Hz, depending on the dynamics (FRAP recovery rates, MSD of microspheres, and fusion of coacervates) of the samples. Imaging data were processed and analyzed using ImageJ.

Temperature-controlled experiments were conducted using a CherryTemp setup (Cherry Biotech, France) and captured using the spinning disk confocal system described above. In each experiment, a freshly prepared solution was deposited onto a passivated cover glass, and a thermalization chip was positioned on top of the slide following the manufacturer’s protocol. The glass slide and the chip were separated by 500 µm liquid silicone spacers. Temperature was varied between 20°C and 45°C with a precision of 0.1°C. After each heating or cooling step, the system was allowed to equilibrate for 2 minutes before acquiring the images.

### M6. FRAP

Fluorescence recovery after photobleaching (FRAP) measurements were conducted using a Micro Point pulsed nitrogen-pumped dye laser (405 nm) from Andor Technologies, integrated with the spinning disk confocal microscope. For each experiment, a circular spot with a diameter of ∼1 µm at the center of the complex coacervates was bleached. For experiments involving partial photobleaching, we selected droplets with a minimum diameter of ∼10 µm. Fluorescence recovery was recorded at different frame rates for the various patchyDNA sequences in order to account for their different mobility and to minimize the effects of photobleaching during imaging. For rapidly recovering T-90, GC, and GATC coacervates, movies were acquired at 10 fps (frames per second) (30 s total), while GGCC, GAATTC, and GGATCC coacervates were imaged at 1 fps (300 s total). For GGGCCC and GGAATTCC coacervates, we used 0.1 fps frame rate (30 min total). Temperature-dependent FRAP experiments were conducted using the CherryTemp setup described above.

To determine the characteristic recovery time, the mean fluorescence intensity of the circular bleached region (B(t), 1 µm diameter) and mean intensity at the center of an unbleached coacervate (U(t), 1 µm diameter) were monitored before and after photobleaching. The background noise (N(t)) during this imaging was measured by monitoring a circular area (1 µm diameter) in the solution phase, outside of the droplet. The normalized intensity at the bleach spot, I_N_(t), was calculated using the following relation:

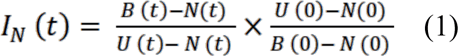

Here, U(0), B(0), and N(0) are mean pre-bleach intensities averaged over 5 frames. The recovery data were double-normalized to account for lost signal due to the bleaching pulse and photobleaching during imaging. The characteristic FRAP recovery time, τ_FRAP_, was calculated by fitting I_N_(t) vs. t with the following equation:

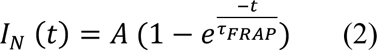

where A is the total recovery fraction. The self-diffusion coefficient (D_self_) of DNA within the coacervates was estimated using the following relation:

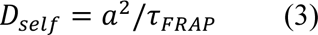

where *a* is the radius of the bleached spot (0.5 µm)

### M7. Condensate fusion

Spontaneous fusion events were manually identified in movies of condensates recorded using either an Andor iXon EM-CCD (for movies at < 20 fps) or an Andor Zyla sCMOS (for movies at > 20 fps). Fusion events for T-90 and GC coacervates were recorded at 200 fps, while coacervates of the other patchyDNA were recorded at rates ranging from 10 to 0.1 fps. The characteristic fusion time, τ_fusion_, was calculated from the temporal evolution of the aspect ratio (AR(t)) of a coacervate after contact using the following relation:

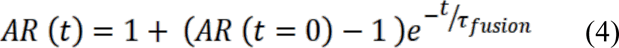

We chose fusion events between droplets of roughly similar size. In a typical fusion event, AR(t=0) ≈ 2 and relaxes to ∼1 over time. For each patchyDNA, we examined fusion events across a range of droplet sizes. The inverse capillary velocity (η/γ) for was then calculated using the following relation:

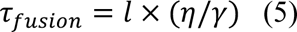

Where *l* is the size of the coacervate after fusion, defined as:

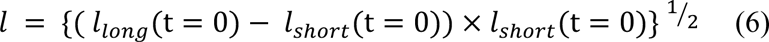

Here, *l*_long_ and *l*_short_ are the long and short axes of the coacervate, respectively. Coacervate fusion experiments were performed on freshly passivated 384-well plates to minimize effects of surface adsorption on fusion dynamics.

### M8. Single particle tracking

The viscosity of coacervates was calculated by tracking trajectories of passivated fluorescent polystyrene beads (diameter = 100 nm, Thermo Fisher, USA) embedded in coacervates. Bead trajectories were imaged using Andor iXon EMCCD with a 561 nm excitation laser at 5 fps and analyzed using the Trackmate plugin in ImageJ^60^.

The diffusion coefficient of beads (D_probe_) was calculated from the mean squared displacement (MSD) using the following relation:

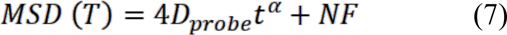

Here, α is the diffusion exponent, and NF is the noise floor, estimated by tracking beads stuck at the bottom of the coacervates. Coacervates viscosity was estimated using the Stokes-Einstein relation:

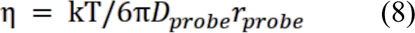

Here, r_probe_ = 50 nm. For all MSD experiments, beads diffusing far away from the coacervate-solution interface were examined in order to minimize any surface effects on bead trajectories. Where indicated, the temperature was modulated using the CherryTemp instrument, as described above, and for each temperature, NF was determined independently.

### M9. FRET

To determine FRET efficiency within complex coacervates, we used a method similar to that was used by Nott et. al.^61^ In brief, three fluorescent images: DD, DA and AA at a fixed z-plane are captured. Here DD = donor emission after donor excitation; DA (FRET signal) = acceptor emission after donor excitation and AA = acceptor emission after acceptor excitation. The FRET efficiency (F) at a given region of interest is then given by,

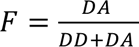

The donor (Cy3) and acceptor (Cy5) fluorophores were incorporated either at the 5’ terminus or internally in the DNA sequence (see Supplementary info for sequences). Cy3 and Cy5 were excited by 561 nm and 637 nm lasers respectively and their emissions were collected using 600-50 nm and 700-75 nm bandpass filters respectively. To correct for spectral overlap, we calculated corrected FRET efficiency by using three sets of samples: sample containing (a) both Cy3-Cy5 (DD_Cy3-Cy5_, DA_Cy3-Cy5_, AA_Cy3-Cy5_), (b) only Cy3 (DD_Cy3_, DA_Cy3_, AA_Cy3_) and (c) only Cy5 (DD_Cy5_, DA_Cy5_, AA_Cy5_). The corrected FRET efficiency, F_corr_ for each pixel within the region of interest was calculated by following the relation,

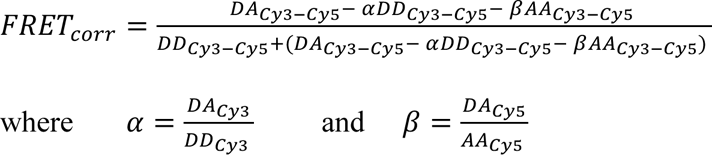

Low intensity values corresponding to DD_Cy5_ and AA_Cy5_ confirmed that signal bleedthrough between channels was minimal. All images were background corrected before processing.

## Author contributions

SM and AJ conceptualized the project and designed the experiments. SM conducted the experiments and analyzed the data. SM and AJ interpreted the data. STC provided passivated beads for microrheology experiments. SM and AJ wrote the manuscript with inputs from all other authors.

## Acknowledgements

We thank Ofer Kimchi, Ella King, Daniel Stein, Alexander Alfredo Alexander-Katz, and members of Jain lab for helpful discussions. This work was supported by grants from the NIH (R35GM151111), David and Lucile Packard Foundation, and the Pew Biomedical Trust.

