## Supplementary text for "Sequence programmable nucleic acid coacervates"

**The PDF file includes:**

Supplementary text
Figs. S1 to S14
Tables S1 to S7

### Supplementary text

#### S1. Design of patchyDNA and patchyRNA

The patchy nucleic acids are designed such that they are primarily single stranded and exhibit multiple reversible self-associative hybridization sites. We used polyT (polyU for RNA) as the single stranded scaffold. These sequences do not form appreciable secondary structure at room temperature. We chose 90-mer DNA or 60-mer RNA for this study. These length scales are readily accessible from commercial vendors using solid phase synthesis. To incorporate self-associative hybridization sites, we incorporated short palindromic patches where DNA/RNA may transiently hybridize. To ensure transient hybridization and rapid reversibility under our experimental condition, we limited the length of the patch to ≤8 nucleotides (≤4 nucleotides for RNA). The patches were designed such that the theoretical predicted hybridization energy (see below) was ≤ 20 $k_bT$. To minimize cooperativity between successive hybridization patches, the patches were separated by at least 15 non-hybridizing bases (≥ 12 nucleotides for RNA). We incorporated 4 hybridization patches per strand. We also experimentally validated that these DNA do not form stable duplexes in the solution phase. Under these design constraints, we can theoretically design ~300 unique self-associative patchyDNA sequences using standard nucleotides. This repertoire may be further expanded using non-palindromic heterotypic associations (as exemplified in Fig. 5) or by using modified nucleotides.

We estimated the theoretical hybridization free energy, ΔG, of self-associating patches using NUPACK (www.nupack.org) under the following conditions: 1 M NaCl, NUPACK model DNA04 (allowing dangle and coaxial stacking), 10 mM DNA patch concentration, and desired temperature. This DNA concentration was chosen as the estimated DNA concentration in the coacervate phase is ~10 mM. For RNA, we used the following conditions: 1 M NaCl, NUPACK model RNA06 (allowing dangle and coaxial stacking), 40 mM RNA patch concentration, and the desired temperature. For heterotypic interactions, we calculated free energy ($\Delta G_{hetero}$) for a pair of primary-seconday patch (eg CTCCTC with GAGGAG or CTCCTC with GAGG) under the following conditons: 1 M NaCl, desired T, NUPACK model DNA04, with 5 mM concentration of each patch.

**S2. DNA concentration in the condensed phase**

We estimated the DNA concentration in the coacervate phase ($C_{dense}$) by doping in a known concentration of fluorescently labeled DNA, and comparing the intensity in the droplet to a standard. In brief, we generated a standard curve of fluorescence intensity of Cy5-labeled DNA in dilute solution in conditions that do not induce complex coacervation (DNA diluted in 10 mM Tris pH 7.0) (Fig. S2). Under identical illumination settings, we measured the intensity of labeled DNA in coacervate-inducing conditions (total input DNA 10 μM, 0.4% Cy5 labeled DNA) and estimated $C_{dense}$ by extrapolating the standard curve to the observed fluorescence intensity. For T-90 coacervates, when the input DNA concentration is 10 μM, $C_{dense}$ was ≈ 10 mM (Fig. S2), and the DNA concentration in the dilute phase ($C_{dilute}$) was ≈ 1 μM, indicating a 4 orders of magnitude enrichment in the coacervate phase. Similar results were observed for the other patchyDNA examined (Fig. S2). These measurements provide an order of magnitude estimate of DNA concentration as the high concentration of labeled DNA in the coacervate phase may result in dye quenching, and the local environment may affect dye quantum yield and brightness.

The DNA concentration in the coacervate phase is comparable to the estimated overlap concentration. For a 90-mer single stranded DNA, the radius of gyration, $R_g$, ≈ 5 nm[34]. The estimated overlap concentration[62] where the chains, on an average, begin to overlap with one another, is given by, $C_{overlap} \sim 1/(4/3\ \pi R_g^3 N_A)$ M. Thus, $C_{overlap}$ for 90-mer patchyDNAs is ~1 mM. Comparing $C_{overlap}$ with the DNA concentration in the dilute and the coacervate phases, we infer that the DNA in solution phase is in the dilute regime ($C_{dilute} << C_{overlap}$) while within coacervates is at the onset or slightly more than the overlap concentration (Fig. S2). Thus, patchyDNA chains in the coacervate phase interact with each other and may form inter-molecular base pairs.

As an alternative approach, we used FRET to experimentally examine the mean inter-molecular distance between DNA chains. PatchyDNAs were labeled at the 5' end with either a fluorescence donor (Cy3) or an acceptor (Cy5) (see Supplementary table 3). 500 nM of donor labeled DNA was mixed with increasing concentration of acceptor labeled DNA (0 – 500 nM), and unlabeled DNA was added to a final total DNA concentration of 10 μM ($C_{input}$ = 10 μM). In the dilute phase, donor signal did not appreciably change as we increased the acceptor concentration (Fig. S3). However, in the coacervate phase, the donor fluorescence progressively diminished with increasing concentration of the acceptor labeled DNA indicative of FRET between the two dyes. This FRET signal indicates that the inter-probe distance in the coacervate phase was comparable to Forster radius, $R_0$. For Cy3-Cy5 dye pair $R_0$ ≈ 5 nm[33], which is comparable to $R_g$ for 90-mer ssDNA (≈ 5 nm)[34]. These observations further indicate that within complex coacervates the DNA density is comparable to the overlap concentration.

### S3. Estimation of critical salt concentration

To determine the critical salt concentration (C*), patchyDNA complex coacervates were formed at NaCl concentrations ranging from 0 -140 mM (see Methods). Freshly prepared coacervate mixture was transferred to a passivated 384 glass-bottom well plate and subsequently imaged using spinning disk confocal microscope. For each patchyDNA, we assembled complex coacervates with increasing NaCl concentration in 10 mM steps (0 mM, 10 mM, 20 mM and so on). C* was determined as the NaCl concentration, at and beyond which no complex coacervates were observed under microscope. As an alternative to our microscopy analysis, we conducted turbidity measurements by examining the absorbance at 400 nm using a Nanodrop 8000 UV-vis Spectrophotometer (Thermofisher, USA). In brief, the coacervate components were thoroughly mixed and equilibrated at room temperature (22°C) for a few minutes. 1 µl of the mixture was transferred onto the spectrophotometer pedal and the absorbance/transmittance between 220 – 800 nm was measured. Turbidity (= 100 – transmittance%) was calculated using the transmittance at 400 nm. Then the turbidity of coacervate mixtures at all NaCl concentrations was normalized using the turbidity at 0 mM NaCl.

To determine the DNA concentration in the supernatant phase, 100 µl of complex coacervate mixture were prepared at a desired NaCl concentration (final concentration 0 – 50 mM) in 1.5 ml tubes. The tube was then centrifuged at 20,000g for 20 min. After the spin, the complex coacervates accumulated at the bottom of the tube. We collected 50 µl of the supernatant in a new tube and again centrifuged it at 12,000g for 10 min. After this second spin, 20 µl of supernatant was collected. DNA concentration in the supernatant was estimated by the absorbance at 260 nm using the theoretical extinction coefficient provided by IDT. Soluble DNA fraction was calculated by measuring the ratio of DNA concentration in the supernatant with DNA concentration in spermine free DNA solution.

**Supplementary figure legends:**

**Fig. S1. Spermine induces complex coacervation of patchyDNA**. (A) Schematics showing that when DNA (T-90 or patchyDNA) is mixed with a polycation, the mixture phase separates and forms a DNA dense complex coacervate phase and a DNA dilute, solution phase. (B-E) T-90/spermine coacervates behave like liquids. (B) Representative fluorescent micrographs showing that non-base pairing T-90 DNA forms spherical complex coacervates in the presence of spermine (10 μM DNA, 4 mM spermine, 10 mM Tris pH 7.0, at T = 22°C). Scale bar, 10 μm. Representative fluorescence micrographs (C) and corresponding recovery curve (D) showing rapid recovery of fluorescence intensity after partial photobleaching of a T-90 coacervate. The red arrow depicts the bleached spot. Scale bar in (C), 8 μm. Data points in (D) represent mean ± SD, n = 5 coacervates. (E) T-90 coacervates coalesce and rapidly relax to a spherical shape. Representative fluorescence micrographs showing coalescence of two T-90/spermine condensates. (F) Representative micrographs of patchyDNA/spermine coacervates for patchyDNA with the indicated patch sequences (10 μM DNA, 4 mM spermine, 10 mM Tris pH 7.0). Scale bars in E, F denote 10 μm.

**Fig. S2. Estimation of DNA concentration in complex coacervates.** (A) Schematic showing three different concentration regimes of a polymer solution. Left panel: dilute solution, where polymer concentration ($C_{polymer}$) is lower than the overlap concentration ($C_{overlap}$) and the polymer chains remain far from each other, limiting inter-chain interactions. When $C_{polymer}$ is comparable to (middle panel) or higher (right panel) than $C_{overlap}$, inter-chain interactions become likely. (B) Standard curve for calibration of fluorescence intensity as a function of Cy5 labeled DNA concentration (resuspended in 10 mM Tris pH 7.0, T= 22°C). Each data point denotes mean ± SD, n ≥15 imaging areas. The lines denote linear fits with the following slopes ± SE: T-90 = 592 ± 3;

GC = 745 ± 3; GGCC = 661 ± 3; GGATCC = 597 ± 3; GGAATTCC 681 ± 4. We measured DNA concentration in the DNA-dense phase ($C_{dense}$) and the DNA-poor ($C_{dilute}$) phase. (C) Plot showing $C_{dense}$ (gray bars) and $C_{dilute}$ (black bars) for the indicated patchyDNA. $C_{dense}$ was ~ 1- 10 mM and $C_{dilute} \approx 1$ μM when the input DNA concentration is 10 μM, in a buffer with 4 mM spermine, 10 mM Tris pH 7.0, at T= 22°C.

**Fig. S3. FRET between DNA molecules in the coacervate phase.** (A) Fluorescence micrographs showing titration of FRET acceptor probes (Cy5-DNA) in dilute DNA solution (left, total DNA = 10 μM, Tris pH 7.0 = 10 mM, NaCl = 0 mM, T = 22°C) and within DNA/spermine coacervates (right, total DNA = 10 μM, spermine = 4 mM, Tris 7.0 = 10 mM, NaCl = 0 mM, T = 22°C) for T-90 and GGATCC patchyDNA. In all experiments 500 nM of FRET donor probe (Cy3-DNA) was used. In dilute solution, the donor emission upon donor excitation remains unchanged with- and without- the acceptor labeled strand. Within coacervates, donor emission gradually decreased with the molar fraction of acceptor (A, right panel). The donor and acceptor probes for T-90 and GGATCC are listed in the Supporting Table 3. Scale bars denote 25 μm. (B) Quantification of FRET efficiency in DNA solution (open symbols) and coacervate phase (close symbols) for the indicated patchyDNA.

**Fig. S4. Base-pairing interactions stabilize patchyDNA/spermine complex coacervates against dissolution by NaCl.** (A) Representative fluorescence micrographs for DNA/spermine complex coacervates (10 μM DNA, 4 mM spermine in 10 mM Tris pH 7.0 at T= 22°C) at the indicated NaCl concentrations. Increasing NaCl concentration dissolves DNA/spermine complex coacervates. The critical NaCl concentration required for the dissolution of DNA/spermine coacervates increases with the patch hybridization energy, ε. (B) Phase diagram corresponding to

(A). Scale bars denote 10 μm. (C, D) The critical salt concentration can also be estimated by measuring solution turbidity (absorbance at 400 nm) or by measuring the fraction of DNA in the solution phase obtained by sedimenting the condensates to the bottom of the tube by centrifugation and measuring the DNA concentration in the supernatant. Representative plots showing measured values of turbidity (n = 3, mean ± range) and soluble DNA fraction (n = 3, mean ± range) for T-90/spermine (C), and GGAATTCC/spermine (D) coacervates as a function of NaCl concentration. The black dotted lines indicate the critical salt concentration above which the turbidity is substantially decreased and majority of the DNA is in the dilute phase. The critical salt concentration estimated from the turbidity/soluble DNA fraction measurements agreed with those obtained from the examination of fluorescence micrographs.

**Fig. S5. FRAP curves of patchyDNA/spermine coacervates.** (A) Fluorescence recovery curves for the various patchyDNA/spermine complex coacervates with the indicated patch sequence. Each data point is mean ± SD over n ≥ 5 droplets. Characteristic recovery times ($\tau_{FRAP}$) is obtained by fitting the recovery curves to an exponential function. (B) Plot of $\tau_{FRAP}$ within complex coacervates for the various patchyDNAs at room temperature as a function of theoretical estimated patch hybridization energy (ε) at T=22°C. Each data point denotes mean ± SD, n > 15 measurements. (C) Similar to (B) but plotting $\log(D_{self})$ versus ε. The line in (C) is a linear fit of $\log(D_{self})$ with ε (slope = -0.14 ± 0.01, mean ± SE, $R^2$ =0.95).

**Fig. S6. Temperature dependent FRAP recovery of DNA/spermine complex coacervate.** (A) Temperature modestly affects DNA dynamics in T-90/spermine coacervates. FRAP recovery curves at the indicated temperatures. Each data point represents mean ± sd, n = 5 droplets. (B)

Similar to (A) for GGCC/spermine coacervates. The color bar in B shows the range of hybridization energy (ε) for the GGCC interaction patch as temperature is varied between 20-45°C. (C, D) Plots showing logarithm of DNA self-diffusion coefficient (log($D_{self}$), in GGCC/spermine (C) and GGATCC/spermine (D) coacervates as a function of ε. Each data point is mean ± SD, n = 5 droplets. The straight lines in C and D are linear fits of log ($D_{self}$) with ε (slope = -0.17 ± 0.01, $R^2$ =0.88 and -0.14 ± 0.01, $R^2$ =0.9 respectively). The experiments were performed at the following condition: 10 μM DNA, 4 mM spermine, 10 mM Tris pH 7.0.

**Fig. S7. Temperature modestly affects complex coacervation of non-base pairing T-90 DNA.** Representative fluorescent micrographs (n = 3 independent experiments) showing that phase-behavior of T-90/spermine coacervates at the indicated salt concentrations and temperatures.

**Fig. S8. Characterization of passivated beads.** Representative fluorescence micrographs showing that passivation with PEG minimizes labeled DNA adsorption on the bead surface. The stock bead suspension was diluted into buffer alone (10 mM Tris pH7.0, top row) or into 2 μM of 6-FAM labeled patchyDNA, GGATCC (bottom row) and incubated overnight. Scale bars denote 5 μm.

**Fig. S9. Noise floor in particle tracking microrheology.** Noise floor (NF) in bead tracking experiments was measured as the mean square displacement (MSD) of passivated beads stuck at the bottom of the patchyDNA/spermine complex coacervates. (A) Plot showing NF (mean of n = 5 trajectories) at room temperature (22°C) and at the indicated temperatures (25- 45°) (B). To minimize the impact of errors due to experimental/measurement limitations, we calculated viscosity of coacervates where MSD was at least an order of magnitude higher than the NF. For instance, at room temperature we limited our analysis to DNA with $\varepsilon \leq 14$.

**Fig. S10. Temperature dependence of viscosity of DNA/spermine coacervates.** (A, B) Mean squared displacement of 100 nm diameter passivated beads in T-90/spermine (A) and GGATCC/spermine (B) complex coacervates at the indicated temperatures. The color bar in B shows the hybridization energy, ε, for patch sequence GGATCC between 20-45°C. (C, D) The bead displacement can be used to estimate viscosity (η). Plots showing η for coacervates of patchyDNA with patch sequence GGCC (C) and GGATCC (D) as a function of temperature.

**Fig. S11. Coalescence rates of patchyDNA/spermine coacervates.** (A) Plots showing the characteristic time for fusion, $\tau_{fusion}$, for the indicated patchyDNA. Each data point denotes mean ± range of $\tau_{fusion}$ obtained from the exponential fit of aspect ratio with time for a fusion event. The slope of the line provides an estimate of the inverse capillary velocity, $\eta/\gamma$. (B) Plot showing interfacial tension, γ, of DNA/spermine complex coacervates modestly varies with the hybridization energy, ε. γ was calculated by dividing η by $\eta/\gamma$ of respective patchyDNA. $\eta/\gamma$ values (mean) were obtained from (A) and η was estimated (mean ± SD) from particle tracking experiments.

**Fig. S12. Complex coacervates of DNA with heterotypic cross-links.** (A) Schematics showing the protocol for generating coacervates of DNA mixtures. We mixed primary patchyDNA with secondary patchyDNA at desired molar ratio and added spermine to generate coacervates. For all such experiments, the total DNA concentration was kept constant at 10 μM (4 mM spermine and

10 mM Tris pH 7.0). (B) Plot showing that FRAP recovery rates of T-90 DNA and primary patchyDNA with CTCCTC patches are similar (10 μM DNA, with 4 mM spermine and 10 mM Tris pH 7.0 at T = 22°C). Each data point depicts mean ± SD (n ≥ 5). (C) Plot showing characteristic recovery time, $\tau_{FRAP}$, of primary patchyDNA with patch sequence CTCCTC mixed at indicated molar ratios of the secondary cross-linking strand with the patch sequence GAGGAG. Each data point depicts mean ± SD (n ≥ 3).

**Fig. S13. Coalescence rates of patchyRNA/poly-lysine condensates.** Plots showing the characteristic time for fusion, $\tau_{fusion}$, for the indicated patchyRNA/poly-lysine coacervates mixing 16 μM RNA with 0.5 mM poly-lysine in 10 mM Tris pH 7.0 at T= 22°C. Each data point denotes mean ± range of $\tau_{fusion}$ obtained from the exponential fit of aspect ratio with time for a fusion event. The slope of the line provides an estimate of the inverse capillary velocity, $\eta/\gamma$. (B) Plot showing interfacial tension, $\gamma$, of RNA/poly-lysine complex coacervates modestly varies with the hybridization energy, $\varepsilon_{RNA}$. The values were calculated using $\eta/\gamma$ measurements from (A) and $\eta$ from single particle tracking experiments.

**Fig. S14. Temperature dependent material properties of patchyRNA/poly-lysine condensates.** MSD tracks of 100 nm diameter beads (A, C) and calculated viscosity (B, D, E) of RNA without self-associative patches r(U-60) (A, B), with patch sequence r(GC) (C, D) and r(GAUC) (E). The hybridization energy is varied by changing the temperature. The color bars denote theoretical predicted hybridization energy ($\varepsilon_{RNA}$) in the temperature range 25-45°C. MSD data are mean of $n \geq 5$ experiments. Viscosity measurements denote mean ± SD across $n \geq 5$ independent experiments.

**Table S1.**

List of patchyDNA, used fluorescent tags (for FRAP) and self-binding interactions of patches at room temperature

| DNA name | 5' - 3' sequence | Fluorescent tag | patch sequence | length of patches (nt) | %GC in patches | -ΔG (kcal/mol) at T=295 K | ε= -ΔG/RT at T = 295 K |
|---|---|---|---|---|---|---|---|
| T-90 | TTTTTTTTTTTTTTTTTTTTTTTTTTTTTTTTTTTTTTTTTTTTT<br>TTTTTTTTTTTTTT TTTTTTTTTTTTTTTTTTTTTTT TTTTTTTT | 5’-Cy5 | - | 0 | 0 | 0 | 0.00 |
| GC | TTTTTTTTTGCTTTTTTTTTTTTTTTTTTTTGCTTTTTTTTTT<br>TTTTTTTTTTGCTT TTTTTTTTTTTTTTTTTTTGCTT<br>TTTTTTTT | 5’-Cy5 | GC | 2 | 100 | 3.09 | 5.26 |
| GTAC | TTTTTTTTGTACTTTTTTTTTTTTTTTTTT<br>TGTACTTTTTTTTTTTTTTTTTT<br>TGTACTTTTTTTTTTTTTTTTTT TGTACTTTTTTTTT | 5’-Cy5 | GTAC | 4 | 50 | 4.93 | 8.40 |
| GGCC | TTTTTTTTGGCCTTTTTTTTTTTTTTTTTT<br>TGGCCTTTTTTTTTTTTTTTTTT<br>TGGCCTTTTTTTTTTTTTTTTTT TGGCCTTTTTTTTT | 5’-Cy5 | GGCC | 4 | 100 | 7.36 | 12.54 |
| CGCG | TTTTTTTTCGCGTTTTTTTTTTTTTTTTTT<br>TCGCGTTTTTTTTTTTTTTTTTT<br>TCGCGTTTTTTTTTTTTTTTTTT TCGCGTTTTTTTTT | | CGCG | 4 | 100 | 8.24 | 14.04 |
| GCGC | TTTTTTTTGCGCTTTTTTTTTTTTTTTTTT<br>TGCGCTTTTTTTTTTTTTTTTTT<br>TGCGCTTTTTTTTTTTTTTTTTT TGCGCTTTTTTTTT | | GCGC | 4 | 100 | 8.27 | 14.09 |
| GAATTC | TTTTTTTGAATTCTTTTTTTTTTTTTTTTT<br>GAATTCTTTTTTTTTTTTTTTTT<br>GAATTCTTTTTTTTTTTTTTTTT GAATTCTTTTTTTT | 5’-Cy5 | GAATTC | 6 | 33 | 7.6 | 12.95 |
| GTTAAC | TTTTTTTGTTAACTTTTTTTTTTTTTTTTT<br>GTTAACTTTTTTTTTTTTTTTTT<br>GTTAACTTTTTTTTTTTTTTTTT GTTAACTTTTTTTT | | GTTAAC | 6 | 33 | 7.6 | 12.95 |
| CCTAGG | TTTTTTTCCTAGGTTTTTTTTTTTTTTTTT<br>CCTAGGTTTTTTTTTTTTTTTTT<br>CCTAGGTTTTTTTTTTTTTTTTT CCTAGGTTTTTTTT | | CCTAGG | 6 | 67 | 8.85 | 15.08 |
| GGATCC | TTTTTTTGGATCCTTTTTTTTTTTTTTTTT<br>GGATCCTTTTTTTTTTTTTTTTT<br>GGATCCTTTTTTTTTTTTTTTTT GGATCCTTTTTTTT | 5’-Cy5 | GGATCC | 6 | 67 | 9.21 | 15.69 |
| CCATGG | TTTTTTTCCATGGTTTTTTTTTTTTTTTTT<br>CCATGGTTTTTTTTTTTTTTTTT<br>CCATGGTTTTTTTTTTTTTTTTT CCATGGTTTTTTTT | | CCATGG | 6 | 67 | 9.52 | 16.22 |
| CGATCG | TTTTTTTCGATCGTTTTTTTTTTTTTTTTT<br>CGATCGTTTTTTTTTTTTTTTTT<br>CGATCGTTTTTTTTTTTTTTTTT CGATCGTTTTTTTT | | CGATCG | 6 | 67 | 10.09 | 17.19 |
| GGGCCC | TTTTTTTGGGCCCTTTTTTTTTTTTTTTTT<br>GGGCCCTTTTTTTTTTTTTTTTT<br>GGGCCCTTTTTTTTTTTTTTTTT GGGCCCTTTTTTTT | 5’-Cy5 | GGGCCC | 6 | 100 | 11.64 | 19.83 |
| GGAATTCC | TTTTTTGGAATTCCTTTTTTTTTTTTTGGAATTCCTTTTT<br>TTTTTTTT GGAATTCCTTTTTTTTTTTTTTT<br>GGAATTCCTTTTTT | 5’-Cy5 | GGAATTCC | 8 | 50 | 11.88 | 20.24 |

**Table S2.**

Self-binding interaction of DNA patches at different temperatures

| | GGCC | | GGATCC | |
|---|---|---|---|---|
| Temperature (K) | -ΔG (kcal/mol) | ε= -ΔG/RT | -ΔG (kcal/mol) | ε= -ΔG/RT |
| 293 | 7.48 | 12.83 | 9.41 | 16.14 |
| 298 | 7.18 | 12.11 | 8.9 | 15.01 |
| 303 | 6.87 | 11.39 | 8.39 | 13.91 |
| 308 | 6.56 | 10.70 | 7.88 | 12.86 |
| 313 | 6.25 | 10.03 | 7.36 | 11.82 |
| 318 | 5.93 | 9.37 | 6.85 | 10.82 |

**Table S3.**

List of FRET probes:

| Probes | 5' - 3' sequence |
|---|---|
| T-90-Donor | [Cy3]TTTTTTTTTTTTTTTTTTTTTTTTTTTTTT TTTTTTTTTTTTTTTTTTTTTTT TTTTTTTTTTTTTTTTTTTTTTT TTTTTTTTTTTTTT |
| GGATCC-Donor | [Cy3]TTTTTTTGGATCCTTTTTTTTTTTTTTTT GGATCCTTTTTTTTTTTTTTTTT GGATCCTTTTTTTTTTTTTTTT GGATCCTTTTTTTT |
| T-90-Acceptor | Cy5]TTTTTTTTTTTTTTTTTTTTTTTTTTTTTT TTTTTTTTTTTTTTTTTTTTTTT TTTTTTTTTTTTTTTTTTTTTTT TTTTTTTTTTTTTT |
| GGATCC-Acceptor | [Cy5]TTTTTTTGGATCCTTTTTTTTTTTTTTTT GGATCCTTTTTTTTTTTTTTTTT GGATCCTTTTTTTTTTTTTTTT GGATCCTTTTTTTT |

**Table S4.**

List of DNA sequences used for heterotypic binding:

| patch functionality | DNA name | 5' - 3' sequence | patch sequence | $-\Delta G_{hetero}$ (kcal/mol) with CTCCTC at T = 295 K | $\varepsilon_{heterotypic}=-\Delta G_{hetero}/RT$ at T = 295 K |
|---|---|---|---|---|---|
| primary | CTCCTC | TTTTTTTCTCCTCTTTTTTTTTTTTTTTTT CTCCTCTTTTTTTTTTTTTTTTT CTCCTCTTTTTTTTTTTTTTTTT CTCCTCTTTTTTTT | CTCCTC | - | - |
| secondary | GAGGAG | TTTTTTTGAGGAGTTTTTTTTTTTTTTTTT GAGGAGTTTTTTTTTTTTTTTTT GAGGAGTTTTTTTTTTTTTTTTT GAGGAGTTTTTTTT | GAGGAG | 9.08 | 15.47 |
| secondary | GAGG | TTTTTTTTGAGGTTTTTTTTTTTTTTTTTT TGAGGTTTTTTTTTTTTTTTTTT TGAGGTTTTTTTTTTTTTTTTTT TGAGGTTTTTTTTT | GAGG | 6.74 | 11.48 |
| secondary | AGGA | TTTTTTTTAGGATTTTTTTTTTTTTTTTTT TAGGATTTTTTTTTTTTTTTTTT TAGGATTTTTTTTTTTTTTTTTT TAGGATTTTTTTTT | AGGA | 6.36 | 10.83 |
| secondary | GG | TTTTTTTTAGGATTTTTTTTTTTTTTTTTT TAGGATTTTTTTTTTTTTTTTTT TAGGATTTTTTTTTTTTTTTTTT TAGGATTTTTTTTT | GG | 4.4 | 7.5 |

**Table S5.**

Heterotypic interaction strength at different temperatures:

| | CTCCTC-GAGGAG | |
|---|---|---|
| Temperature (K) | $-\Delta G_{hetero}$ (kcal/mol) | $\varepsilon_{heterotypic} = -\Delta G_{hetero}/RT$ |
| 298 | 8.76 | 14.77234 |
| 303 | 8.24 | 13.66615 |
| 308 | 7.72 | 12.59587 |
| 313 | 7.2 | 11.55978 |
| 318 | 6.68 | 10.55628 |

**Table S6.**

List of patchyRNA sequences and self-binding interactions of patches at room temperature

| RNA name | 5' - 3' sequence | patch sequence | length of patches (nt) | %GC in patches | $-\Delta G$ (kcal/mol) at T = 295 K | $\varepsilon_{RNA}=-\Delta G/RT$ at T = 295 K |
|---|---|---|---|---|---|---|
| U-60 | UUUUUUUUUUUUUUUUUUUUUUUUUUUUUUUUUUUUUUUUUUUUUUUUUU UUUUUUUUUU | - | 0 | 0 | 0 | 0.00 |
| r(GC) | UUUUUGCUUUUUUUUUUUUUUGCUUUUUUUUUUUUUUGCUUUUUUUUUUU UUUGCUUUUU | GC | 2 | 100 | 2.24 | 3.82 |
| r(GAUC) | UUUUGAUCUUUUUUUUUUUUGAUCUUUUUUUUUUUUGAUCUUUUUUUUUU UUGAUCUUUU | GAUC | 4 | 50 | 5.56 | 9.47 |
| r(GGCC) | UUUUGGCCUUUUUUUUUUUUGGCCUUUUUUUUUUUUGGCCUUUUUUUUUU UUGGCCUUUU | GGCC | 4 | 100 | 9.82 | 16.73 |

**Table S7.**
Self-binding interaction of RNA patches at different temperatures:

| | r(GC) | | r(GAUC) | |
|---|---|---|---|---|
| Temperature (K) | -ΔG (kcal/mol) | $\varepsilon_{RNA}$ =-ΔG /RT | -ΔG (kcal/mol) | $\varepsilon_{RNA}$ =-ΔG /RT |
| 298 | 2.15 | 3.625632 | 5.3 | 8.937605 |
| 303 | 2 | 3.317026 | 4.88 | 8.093545 |
| 308 | 1.84 | 3.002124 | 4.45 | 7.260572 |
| 313 | 1.69 | 2.713338 | 4.03 | 6.470268 |
| 318 | 1.53 | 2.417831 | 3.6 | 5.689013 |
