## Supplementary figures for "Sequence programmable nucleic acid coacervates"

Figure S1: Spermine induces complex coacervation of patchyDNA

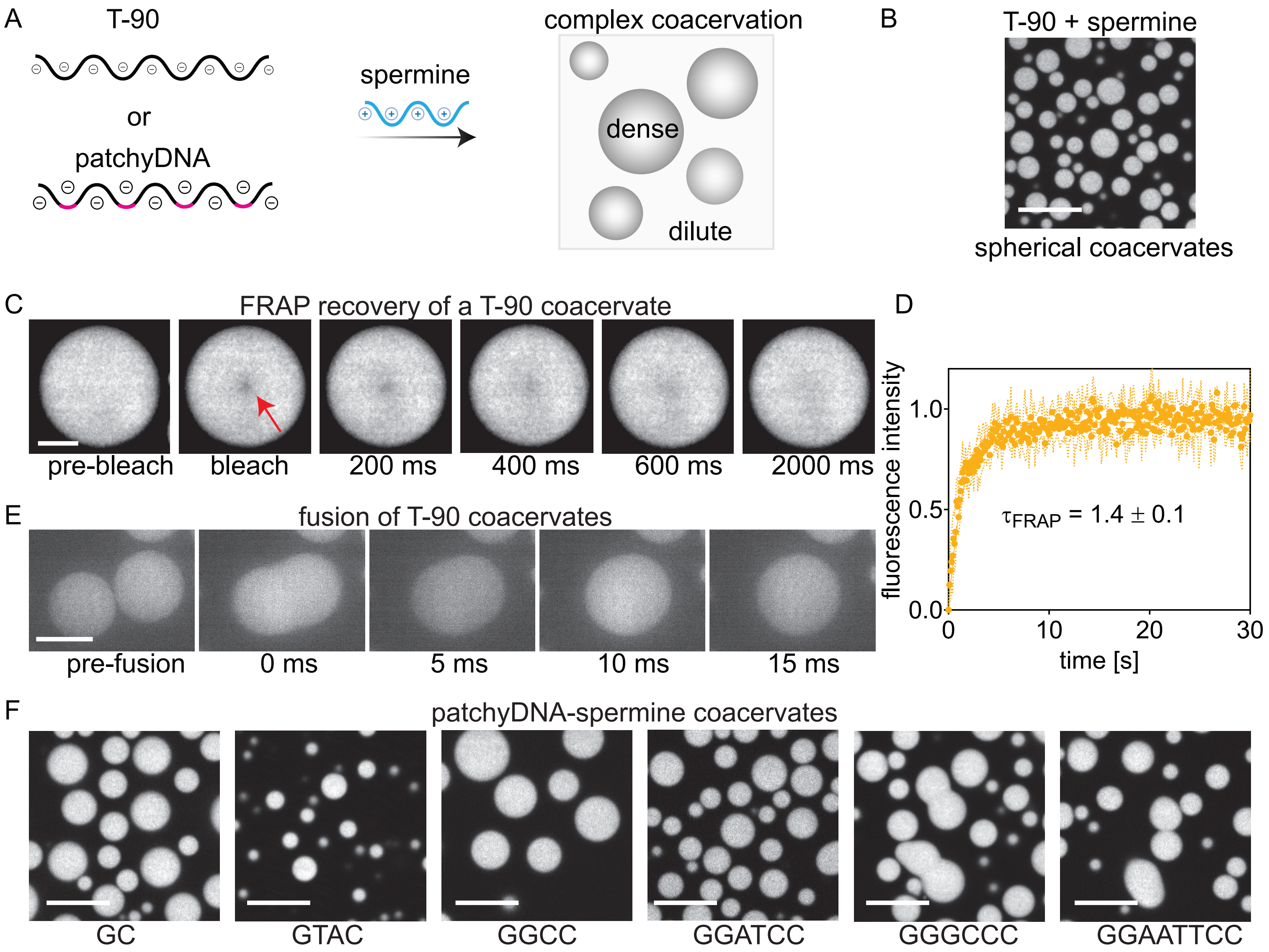

### Figure S2: Estimation of DNA concentration in complex coacervates

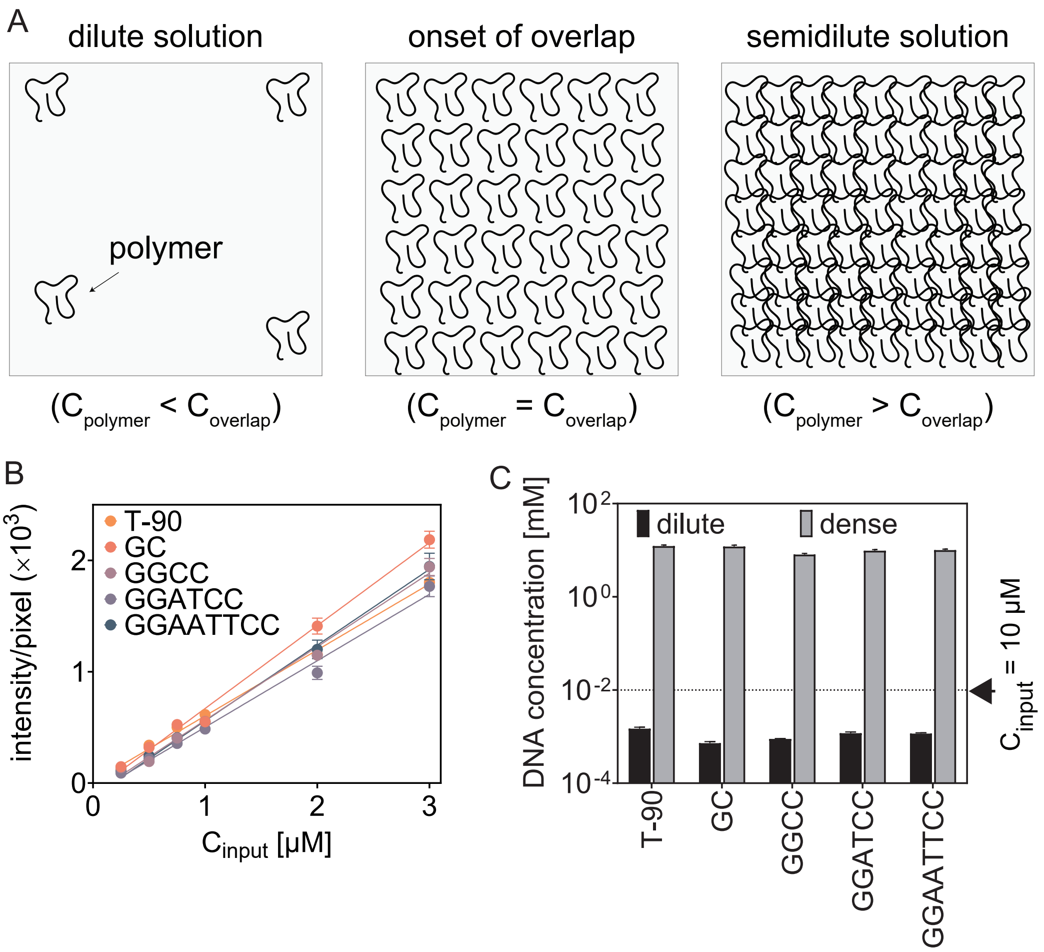

Figure S3: FRET between DNA molecules in the coacervate phase

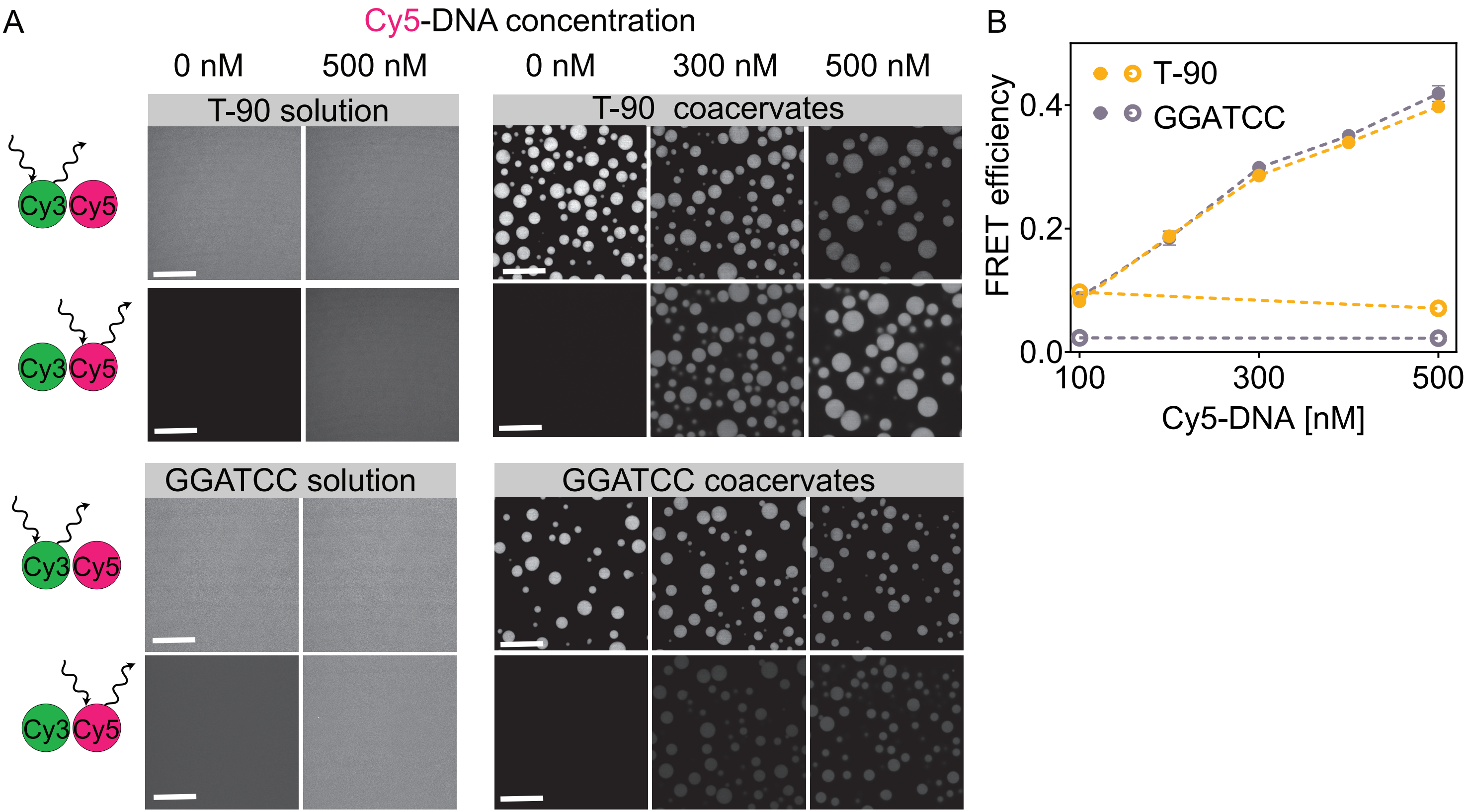

Figure S4: Base-pairing interactions stabilize patchyDNA/spermine complex coacervates against dissolution by NaCl

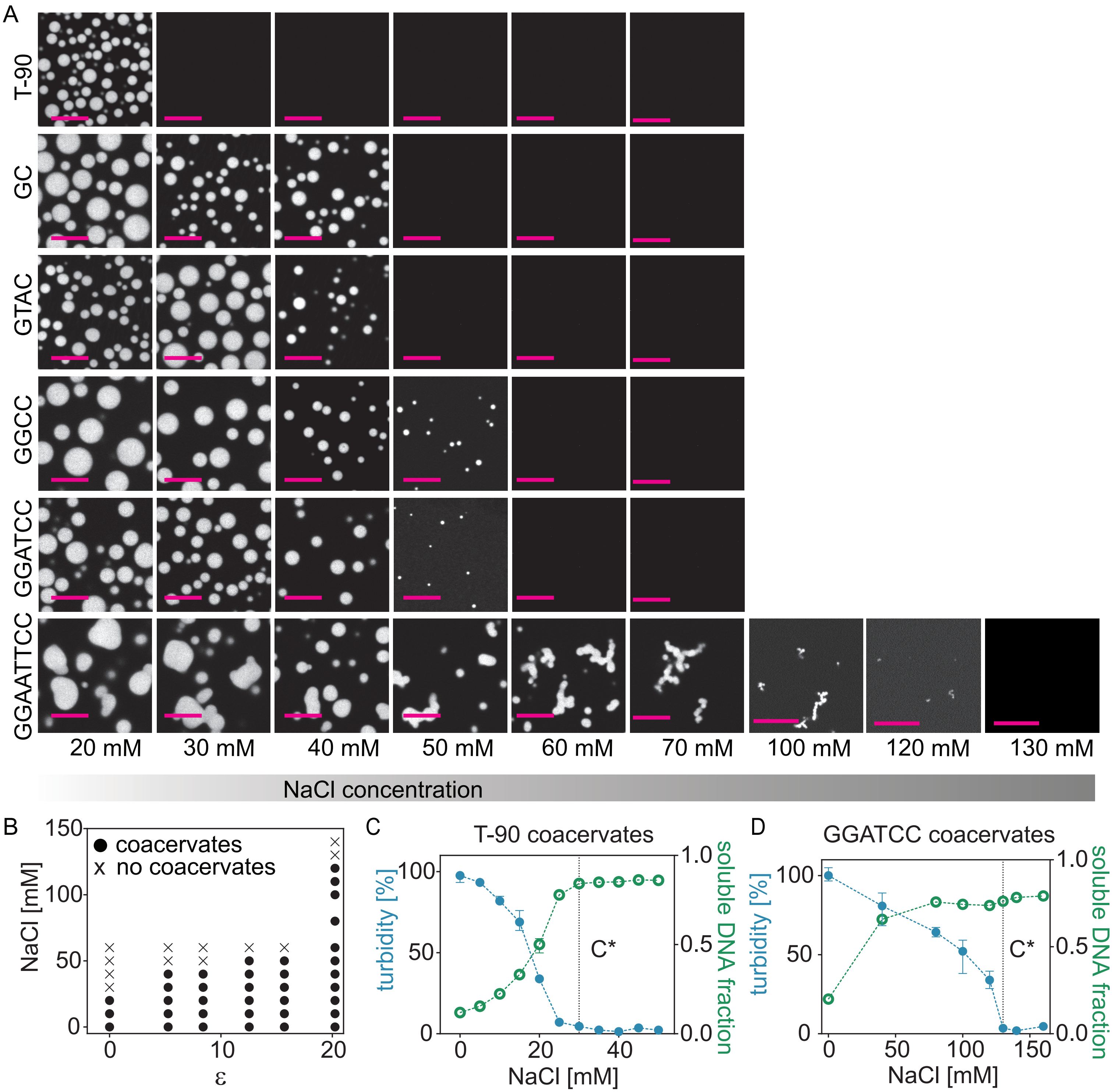

Figure S5: FRAP curves of patchyDNA/spermine complex coacervates

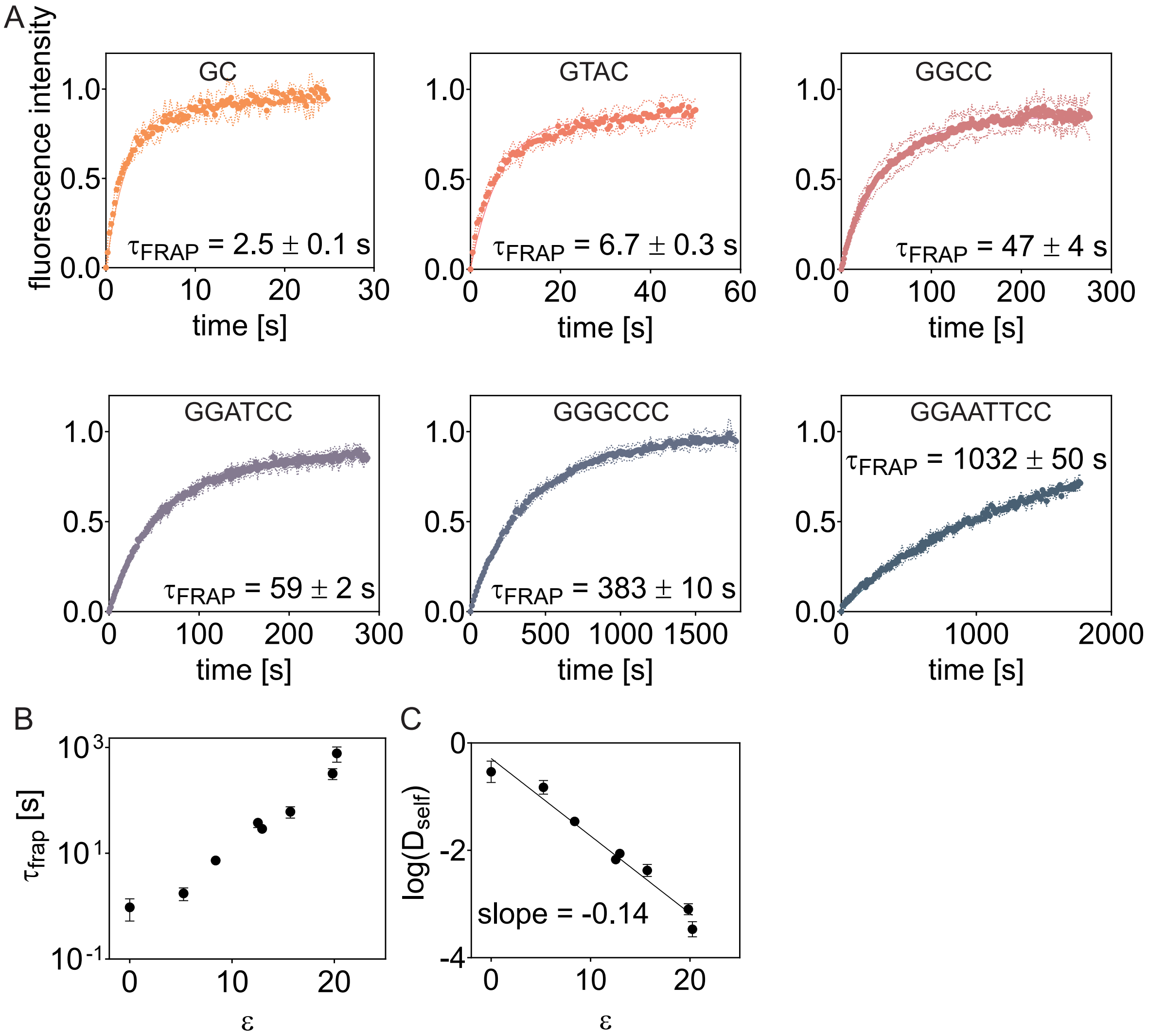

Figure S6: Temperature dependent FRAP recovery of DNA/spermine complex coacervate

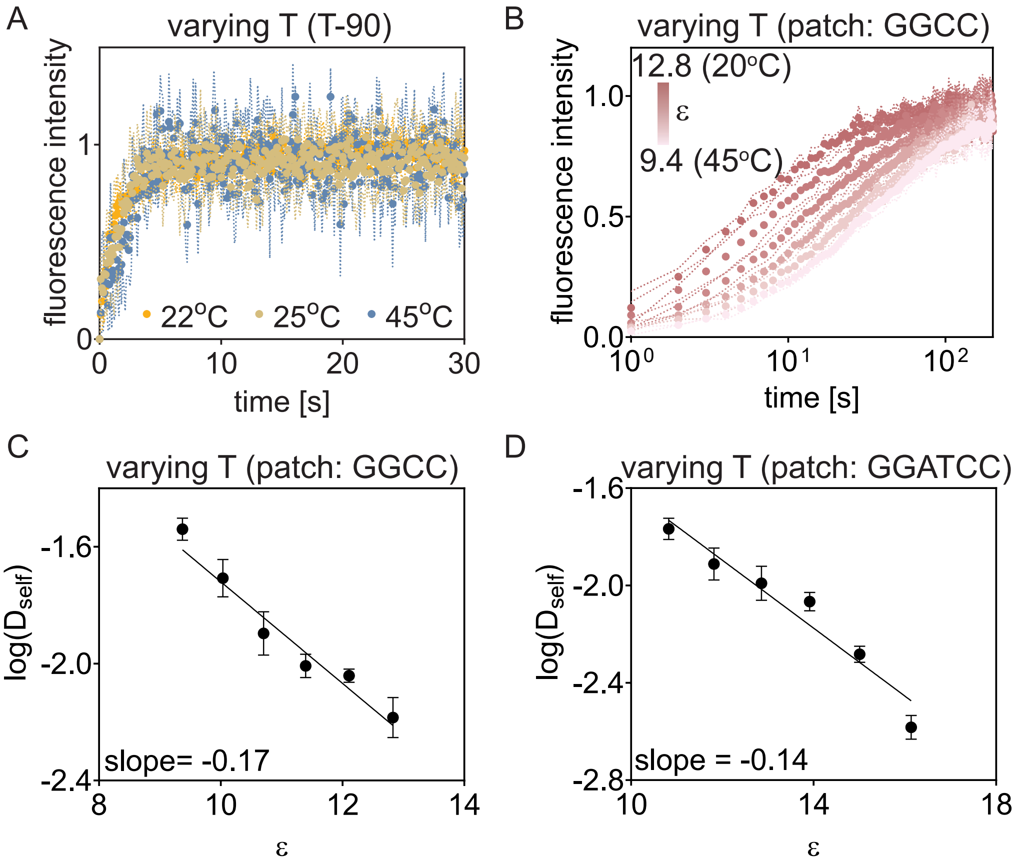

Figure S7: Temperature modestly affects complex coacervation of non-base pairing T-90 DNA

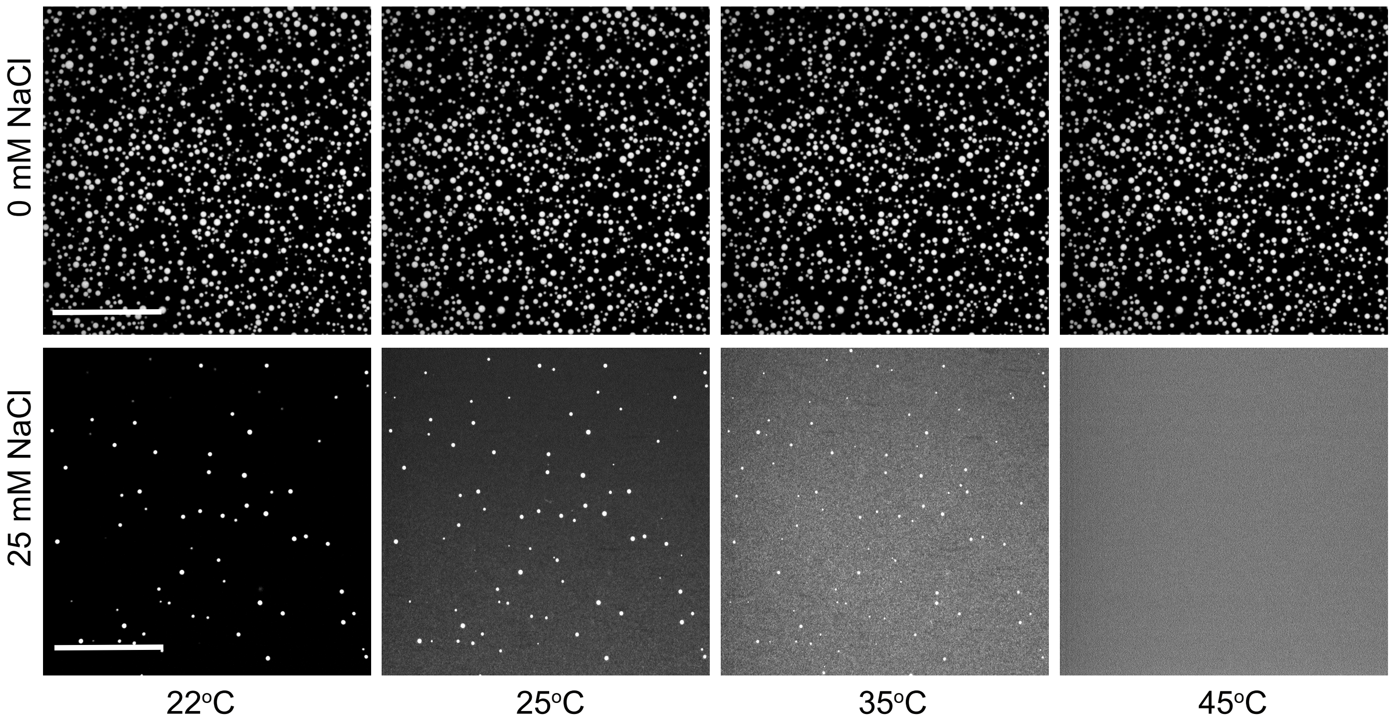

Figure S8: Characterization of passivated beads

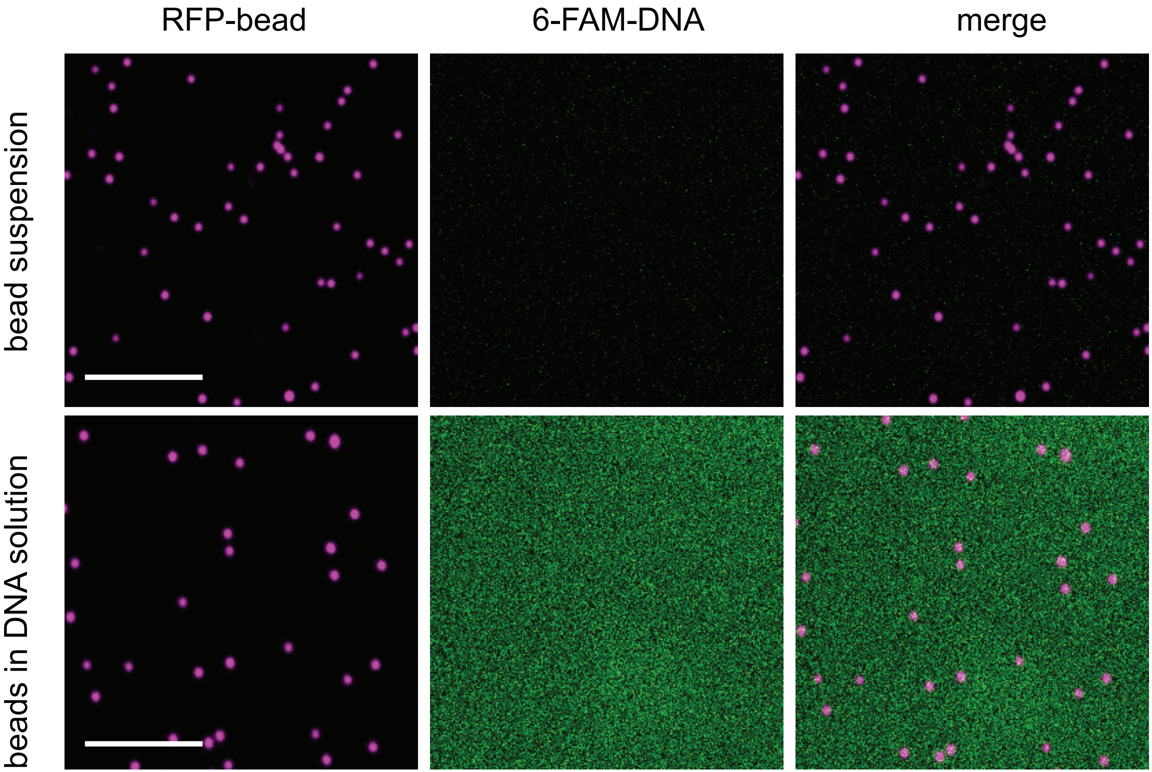

Figure S9: Noise floor in particle tracking microrheology

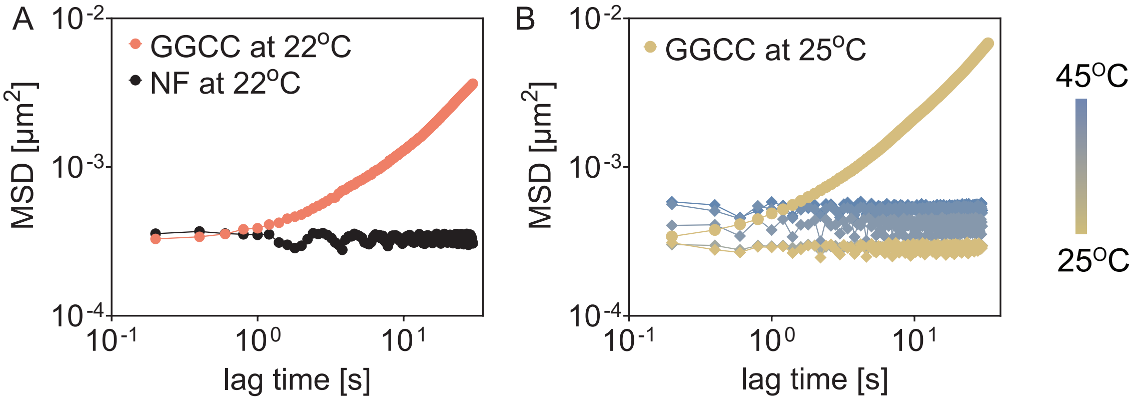

Figure S10: Temperature dependence of viscosity of DNA/spermine complex coacervates

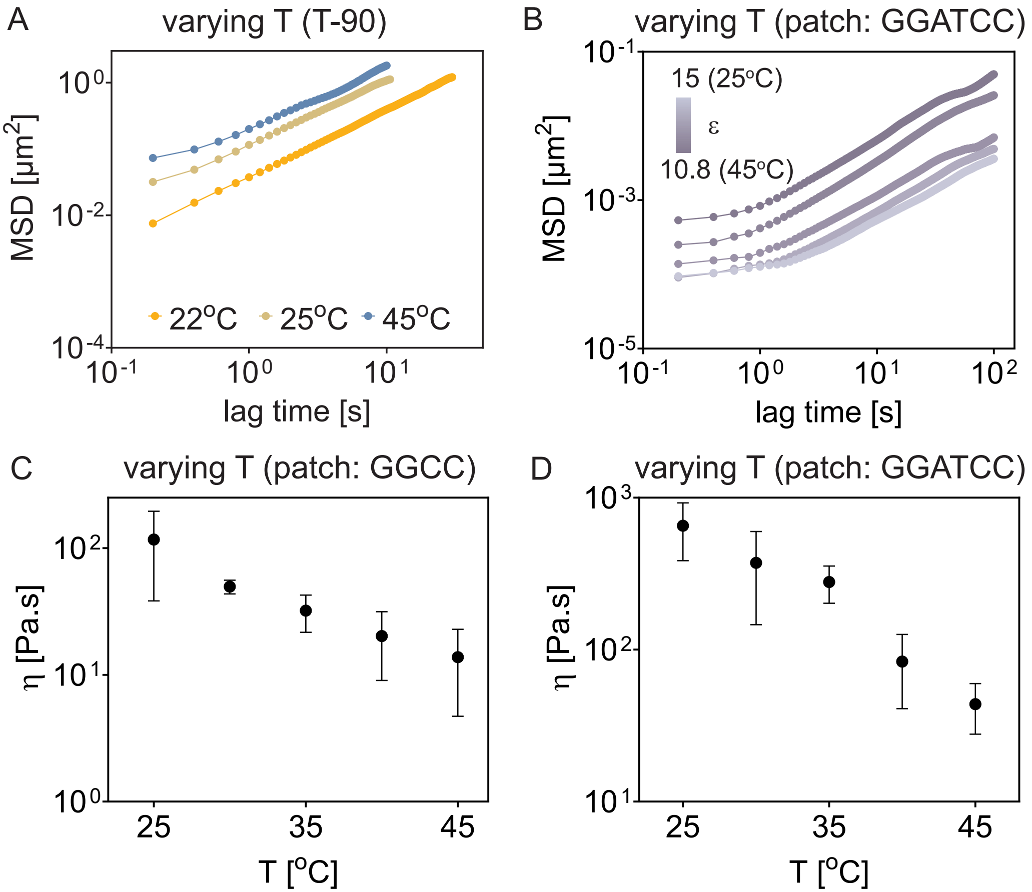

Figure S11: Coalescence rates of patchyDNA/spermine complex coacervates

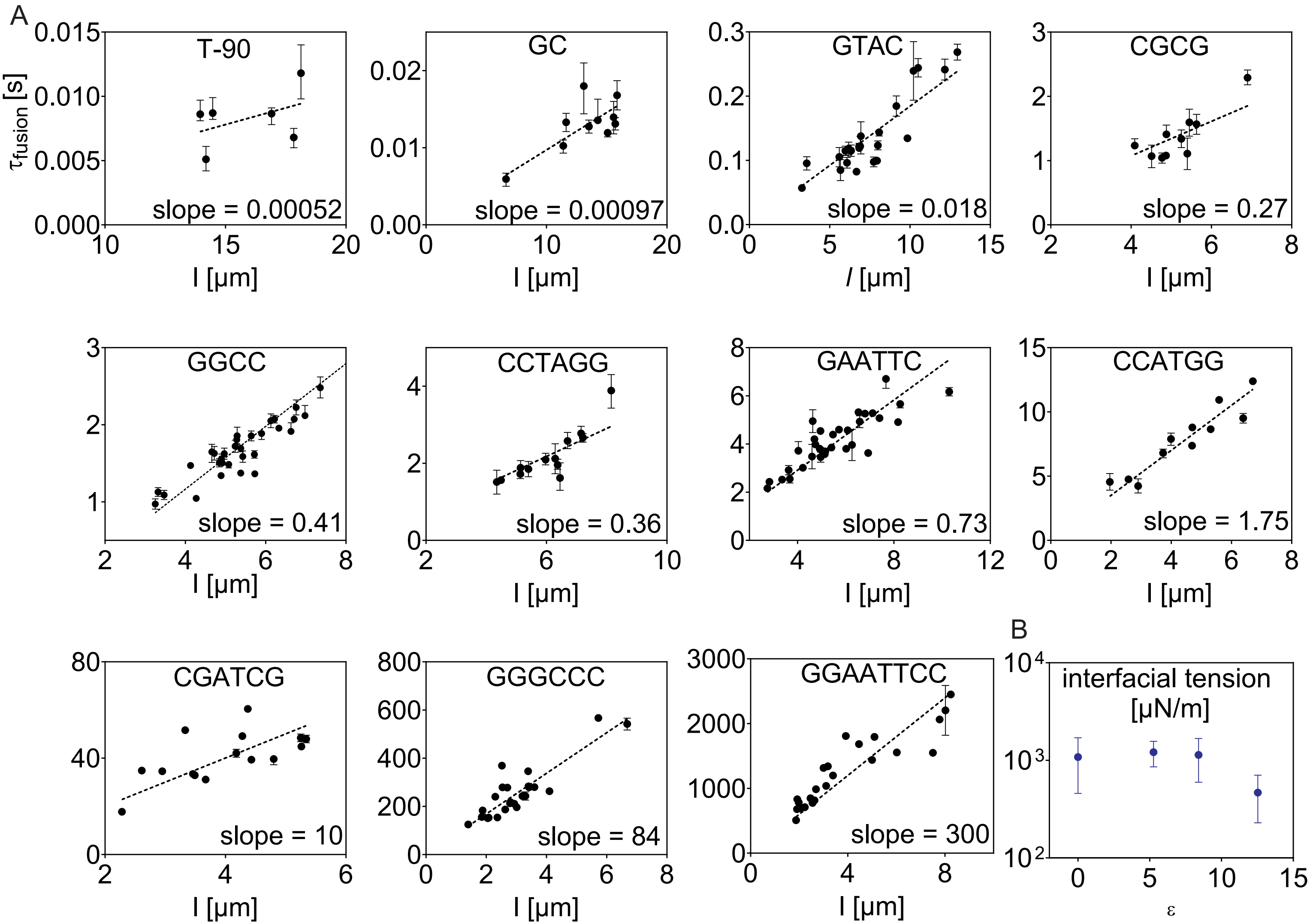

Figure S12: Complex coacervates of DNA with heterotypic cross-links

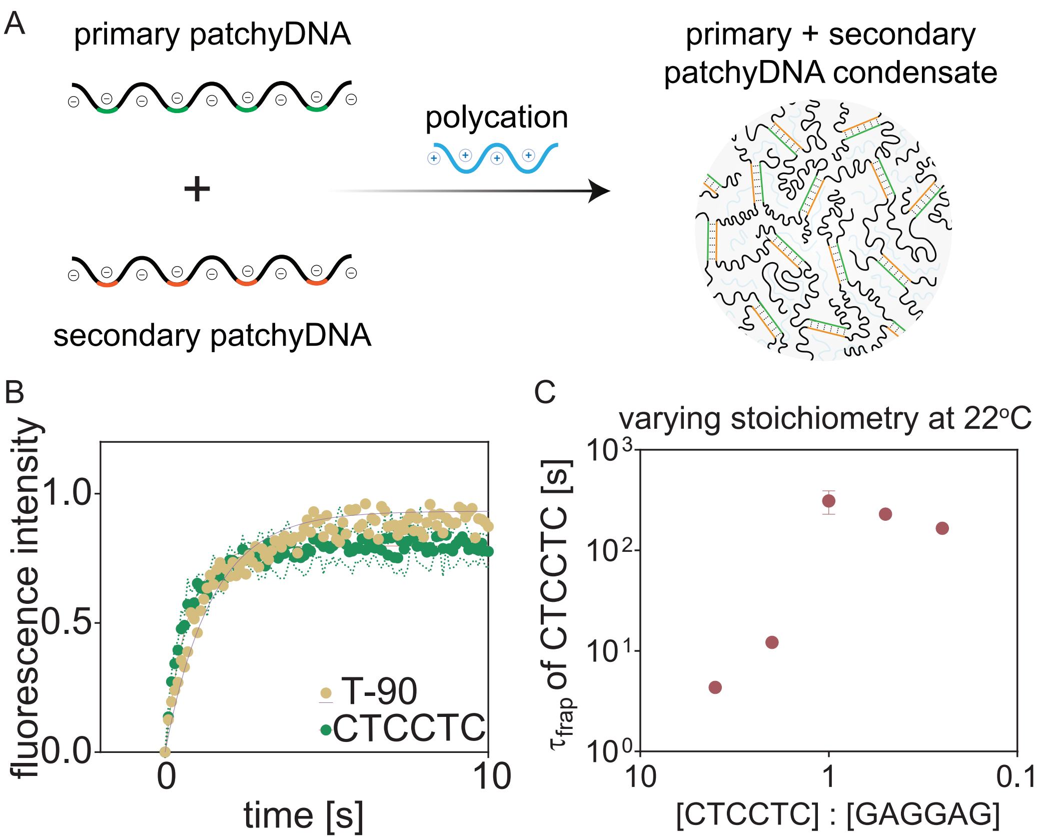

Figure S13: Coalescence rates of patchyRNA/poly-lysine condensates

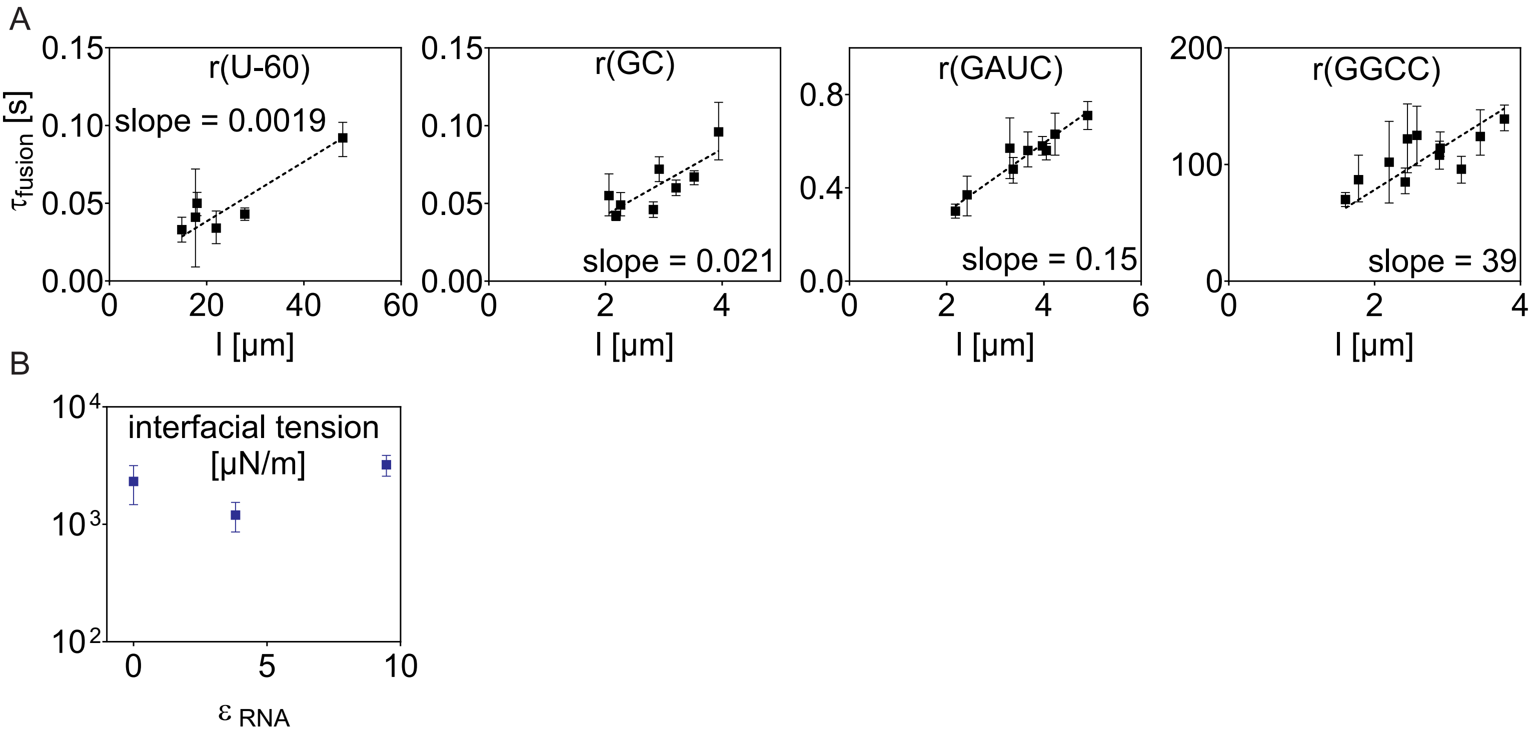

Figure S14: Temperature dependent material properties of patchyRNA/poly-lysine condensates

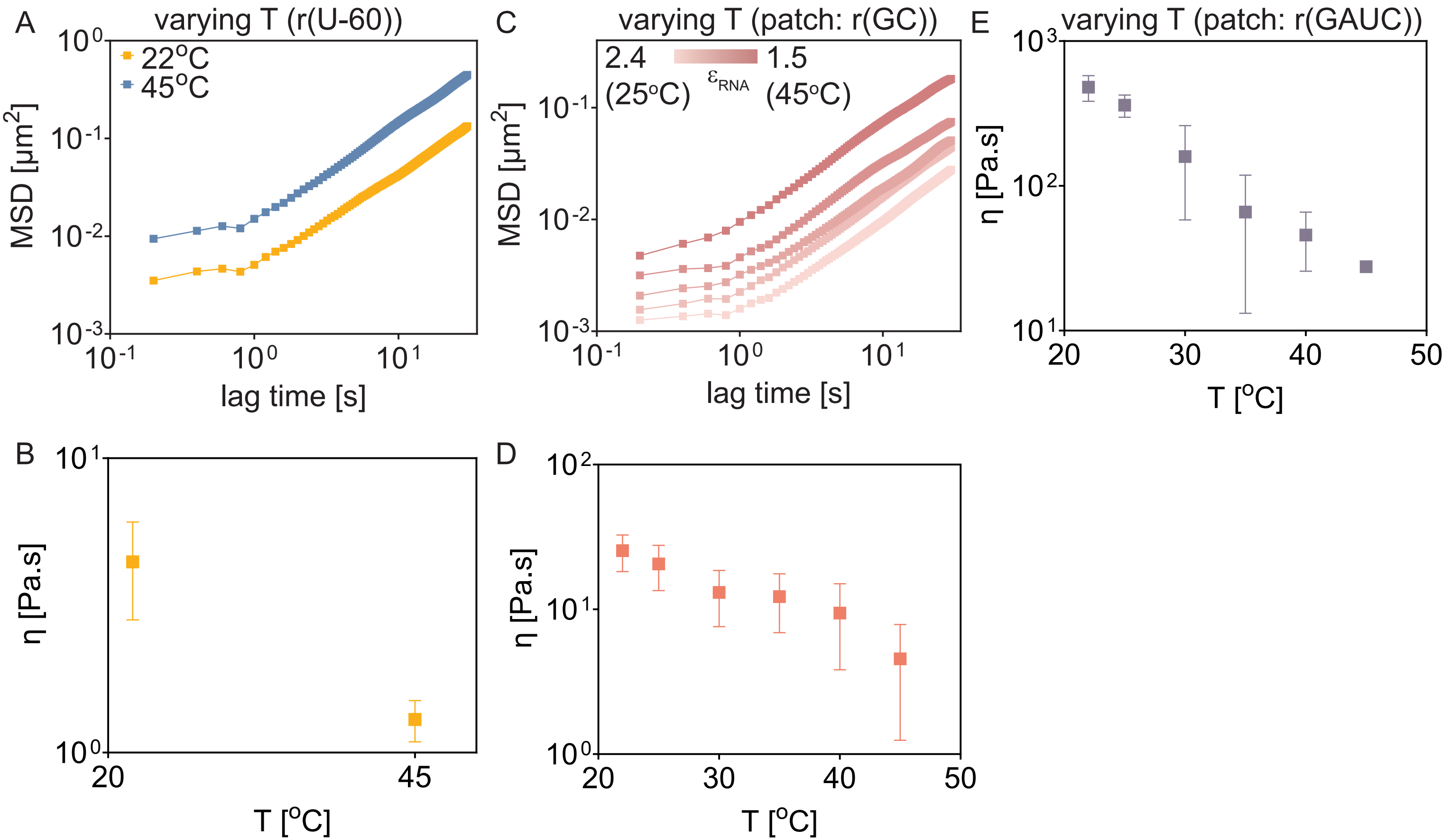
